# BUSTED-PH: Isolating the genomic signatures of convergent phenotypes

**DOI:** 10.64898/2026.01.29.702612

**Authors:** Avery Selberg, Nathan Clark, Anton Nekrutenko, Maria Chikina, Sergei L Kosakovsky Pond

## Abstract

Convergent evolution offers a natural test for adaptive predictability, yet pinpointing its molecular basis remains difficult. Current methods often grapple with distinguishing adaptive convergence from background noise, either by demanding overly stringent identical substitutions or relying on low-resolution evolutionary rate shifts. Here we introduce `BUSTED-PH` (Branch-site Unrestricted Statistical Test for Episodic Diversification – Phenotype), a branch-site codon model that detects phenotype-associated episodic diversifying selection. Unlike standard approaches, `BUSTED-PH` explicitly contrasts selective regimes between phenotype-positive (foreground) and phenotype-negative (background) lineages, effectively winnowing out spurious associations driven by pervasive background adaptation. `BUSTED-PH` has already been applied in numerous independent studies, and here we rigorously validate it using canonical positive controls and simulations to confirm high power and strict false positive control. A genome-wide scan of 120 mammalian species using strict statistical criteria (*FDR* ≤ 0.01) identified 72 genes associated with echolocation; while recovering paradigmatic auditory drivers (e.g., *Prestin, TMC1* ), we also uncover novel candidates in physiological support systems ranging from lipid homeostasis to neural development. A parallel analysis of mammalian gigantism identifies 91 genes linked to musculoskeletal reinforcement, organ size governance, and genomic integrity, characterizing the molecular adaptations required to support massive body size. As projects like Zoonomia and the Vertebrate Genomes Project expand comparative datasets, `BUSTED-PH` provides a robust framework for dissecting the genetic architecture of complex convergent traits.

## Introduction

Convergent evolution, the independent emergence of similar phenotypic traits in distinct lineages, provides a natural experiment for assessing the predictability of evolutionary processes. If phenotypic convergence occurs under similar constraints, it suggests the presence of distinct accessible peaks within the fitness landscape. Canonical systems such as echolocation in mammals [Li et al., 2010, Parker et al., 2013, Liu et al., 2014] and C4 photosynthesis in plants [Christin et al., 2007, Sage, 2004] have served as the crucible for these hypotheses. The advent of the genomic era, propelled by monumental sequencing efforts such as the Zoonomia Project [Zoonomia Consortium, 2020] and the Vertebrate Genomes Project (VGP) [Rhie et al., 2021], has fundamentally shifted the scale of inquiry. The question has evolved from “do phenotypes converge?” to a far more mechanistic and elusive query: “does phenotypic convergence necessitate molecular convergence?” and, conversely, “can we leverage genomic convergence to deconstruct the genetic architecture of complex traits?”

Molecular convergence manifests along a spectrum. At one extreme is *Identical Substitution*, where the same amino acid change occurs in independent lineages, a phenomenon famously debated in the context of the auditory gene *Prestin* in echolocators [Li et al., 2010]. At the other is *Rate Convergence*, characterized by correlated shifts in evolutionary rates, such as acceleration due to positive selection or deceleration due to constraint, without identical mutations [Kowalczyk et al., 2019, Hu et al., 2019]. This rate-based signature is frequently associated with trait loss, such as the degradation of vision genes in subterranean mammals or enhancer loss associated with flightlessness in birds [Sackton et al., 2019]. *Functional Equivalence* represents an intermediate state, where distinct mutations result in similar physicochemical properties [Storz, 2016, Morales et al., 2024]. Detecting these signals requires specific statistical methods. Rate-based methods, including `RERConverge` [Kowalczyk et al., 2019], `PhyloAcc` [ Hu et al., 2019], and `Forward Genomics` [Prudent et al., 2016], effectively identify gene-wide shifts associated with relaxed selection or increased conservation [Partha et al., 2017, Sackton et al., 2019, Kowalczyk et al., 2020]. However, these methods may not resolve adaptive changes restricted to specific sites.

Sequence-based approaches, typically codon models estimating the ratio of nonsynonymous to synonymous substitution rates (*ω* = *dN/dS*), are standard for detecting site-specific positive selection [Kosakovsky Pond and Frost, 2005, Murrell et al., 2015, Wertheim et al., 2015]. However, relating phenotype to genotype is beset by factors such as the “Stokes Shift” in protein evolution, where epistasis and entrenchment reduce the likelihood of identical convergence over time [Pollock et al., 2012]. Additionally, Incomplete Lineage Sorting (ILS) or “hemiplasy” can confound analysis, as mutations on discordant gene trees may be incorrectly inferred as independent convergent events [Mendes and Hahn, 2016]. Standard branch-site codon models test for selection on “foreground” branches but may lack for a rigorous control against background evolution. A gene may be identified as under selection in convergent lineages due to pervasive selection across the phylogeny or model misspecification. Although protocols to filter for foreground-specific selection exist [Kowalczyk et al., 2021], a unified probabilistic framework has not been fully established.

This study presents `BUSTED-PH`, a method that explicitly models and contrasts positive selection across phenotype-defined foreground and background branches. `BUSTED-PH` integrates the `BUSTED-E` framework [Selberg et al., 2025] to account for residual alignment errors and applies a three-part test, each serving a distinct biological rationale:

### 1. Foreground Selection (Test 1)

This tests for evidence of episodic diversifying selection specifically on the lineages possessing the trait (foreground). This is analogous to the alternative hypothesis in standard branch-site models, confirming that adaptation is plausibly occurring in the trait-associated species.

### 2. Background Selection (Test 2)

This tests for positive selection on the background lineages (those lacking the trait). Unlike standard approaches that often assume the background is evolving neutrally or under constraint, this test explicitly checks for “pervasive” selection. A significant result here suggests that the gene is under adaptive pressure across the entire phylogeny, complicating the association with the specific convergent trait.

### 3. Regime Difference (Test 3)

This serves as the crucial contrast, testing whether the distribution of selective pressures (*ω*) significantly differs between foreground and background branches. This distinguishes `BUSTED-PH` from standard models by treating the background not just as a null to be rejected, but as a comparative baseline. Evidence of selection on the foreground is only considered trait-associated if it is distinct from the evolutionary dynamics of the background.

Crucially, an elevated dN/dS ratio on foreground branches indicates distinct selective pressure in these lineages, which may result from convergent adaptation, parallel evolution, or other lineage-specific forces, rather than strictly identical molecular substitutions. Since its integration into the `HyPhy` software package, `BUSTED-PH` has already been implemented in over a dozen published studies to test for trait-associated selection across a variety of biological systems (Table S1; see Table S4 for a comparison with alternative methods). We rigorously validate `BUSTED-PH` using canonical positive controls, genome-wide scans of echolocating mammals and large body size evolution, and simulations to assess power and error rates, acting to disentangle the specific thread of trait-associated adaptation from the knotted skein of genomic history.

## Methods

### Statistical methodology

We model codon evolution using a finite state continuous time Markov model based on the Muse-Gaut family of codon models [Muse and Gaut, 1994]. `BUSTED-PH` builds on the previous `BUSTED` family of models [Murrell et al.,2015, Wisotsky et al., 2020, Lucaci et al., 2023, Selberg et al., 2025], originally developed to test for episodic gene-level positive selection. `BUSTED-PH` employs the same instantaneous rate (*Q*) matrix as in `BUSTED-S` [Wisotsky et al., 2020], where the instantaneous substitution rate between codon *i* and codon *j* is given by:

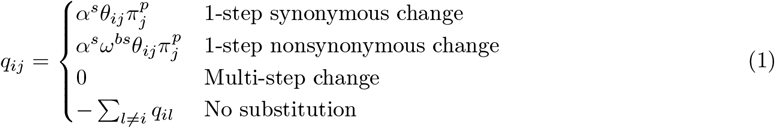

Here, *α*^*s*^ represents the synonymous substitution rate, and *ω*^*bs*^ is the nonsynonymous-to-synonymous rate ratio (*dN/dS*) associated with branch *b* and site class *s*. The parameter *θ*_*ij*_ denotes the nucleotide substitution rate between the codons, and 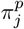 is the position-specific (*p*) equilibrium frequency of the target nucleotide, estimated using the CF3x4 corrected empirical estimator [Pond et al., 2010].

At each branch and site, *ω* is drawn independently from a general discrete distribution with *k* bins (default *k* = 3), allowing selective pressure to vary stochastically across the alignment. Parameters are estimated via maximum likelihood optimization in `HyPhy` [Kosakovsky Pond et al., 2020]. By default, the model includes synonymous rate variation [Wisotsky et al., 2020],; however, users may disable this or enable support for multi-nucleotide substitutions [Lucaci et al., 2023,], via command-line flags.

The `BUSTED-PH` framework explicitly incorporates phenotype data by partitioning branches into *fore-ground* (phenotype-positive) and *background* (phenotype-negative) sets. Distinct *ω* distributions, *S*_*FG*_ and *S*_*BG*_, are then estimated for each partition. Specifically, for any branch in the foreground set, the *ω* at a given site is drawn from *S*_*FG*_ (with bins 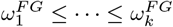), while background branches draw from *S*_*BG*_. The likelihood of the data is computed by marginalizing over all possible assignments of *ω* classes to sites. The number of rate classes, *k*, is a parameter selected a priori (default *k* = 3) based on typical model complexity [Murrell et al., 2015]. However, for smaller datasets or those with limited sequence divergence, this can be reduced (e.g., *k* = 2) to prevent overfitting, as such alignments may lack sufficient information to reliably estimate the parameters of a complex 3-component mixture. If the error sink is enabled, an additional rate class capturing alignment errors, here (*ω*_*e*_ ≥ 1000, weight ≤ 0.25%) is added to both distributions, inferred separately for foreground and background [Selberg et al., 2025].

A critical component of the BUSTED-PH workflow is the assignment of internal branches to the fore-ground or background partitions, given that phenotype data is typically only available for extant species (tips). `HyPhy` implements four distinct strategies for inferring ancestral phenotypes, each corresponding to a specific evolutionary hypothesis:

1. **None (Tips Only):** No internal branches are labeled. This is the most conservative approach, assuming nothing about ancestral states, but may suffer from reduced statistical power due to shorter total branch length, or if evolution took place among the ancestoral lineages.
2. **Conjunctive (All-Descendants):** An internal node is labeled foreground if and only if *all* of its descendants possess the trait. This strategy implies that the trait evolved in the ancestor of a clade and has been largely maintained or finetuned, making it suitable for analyzing trait-defined clades.
3. **Disjunctive (Any-Descendants):** An internal node is labeled foreground if *any* of its descendants possess the trait. This is an inclusive strategy that pushes the origin of the trait deep into the phylogeny, but risks including ancestral branches that did not posess the phenotype.
4. **Parsimony:** Internal branches are labeled based on a maximum parsimony reconstruction of the binary trait. This is appropriate for traits with complex histories of gains and losses, but subject to the limitations of simple basic parsimony.

It is important to note that `BUSTED-PH` does not mandate a specific labeling strategy; users may choose the method best suited to their biological question or provide custom branch labels derived from more sophisticated approaches. For the mammalian analyses in this study, we specifically chose the **Conjunctive** strategy to test for the emergence and finetuning of traits in clades.

### Hypothesis testing and implementation

To test for FG-specific positive selection, we use three likelihood ratio tests (LRT) (Table 1). First, we test for evidence of episodic diversifying selection (EDS) in *FG* branches, comparing model 1 and model 2 with a likelihood ratio test. Second, we test for evidence of positive selection in *BG* branches, with a LRT comparing model 1 and model 3. We derive critical values of the LRT for these two comparisons from a 50:50 mixture model of 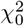 and 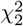 [Self and Liang, 1987, Wisotsky et al., 2020]; this mixture is generally conservative. Third, we test for the difference between *FG* and *BG* branches with a LRT comparing model 1 and model 4 (Table 1, Figure 1). We derive critical values for this final LRT from the 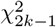 distribution.

**Table 1.** Models run in `BUSTED-PH` to detect FG-specific positive selection. Models 2, 3, and 4 serve as null models, and model 1 serves as the universal alternative model for likelihood ratio test comparisons. Note that if BUSTED-E is enabled, error components are inferred separately for each branch set and are not constrained to be equal across models.

| Model | Explanation | Purpose |
| --- | --- | --- |
| 1 | Independent $\omega$ distributions are estimated for the *FG* and *BG* branches. | Serves as the universal alternative hypothesis for Likelihood Ratio Tests (LRTs) against Models 2, 3, and 4. |
| 2 | The *FG* $\omega$ distribution is constrained to $\omega_k \leq 1$; *BG* remains unconstrained. | Null hypothesis for the test of positive selection on *FG* branches. |
| 3 | The *BG* $\omega$ distribution is constrained to $\omega_k \leq 1$; *FG* remains unconstrained. | Null hypothesis for the test of positive selection on *BG* branches. |
| 4 | A single $\omega$ distribution is estimated for the entire tree, sharing parameters across *FG* and *BG* branches. | Null hypothesis for the test of selective regime differences between *FG* and *BG* branches. |

**Figure 1.**
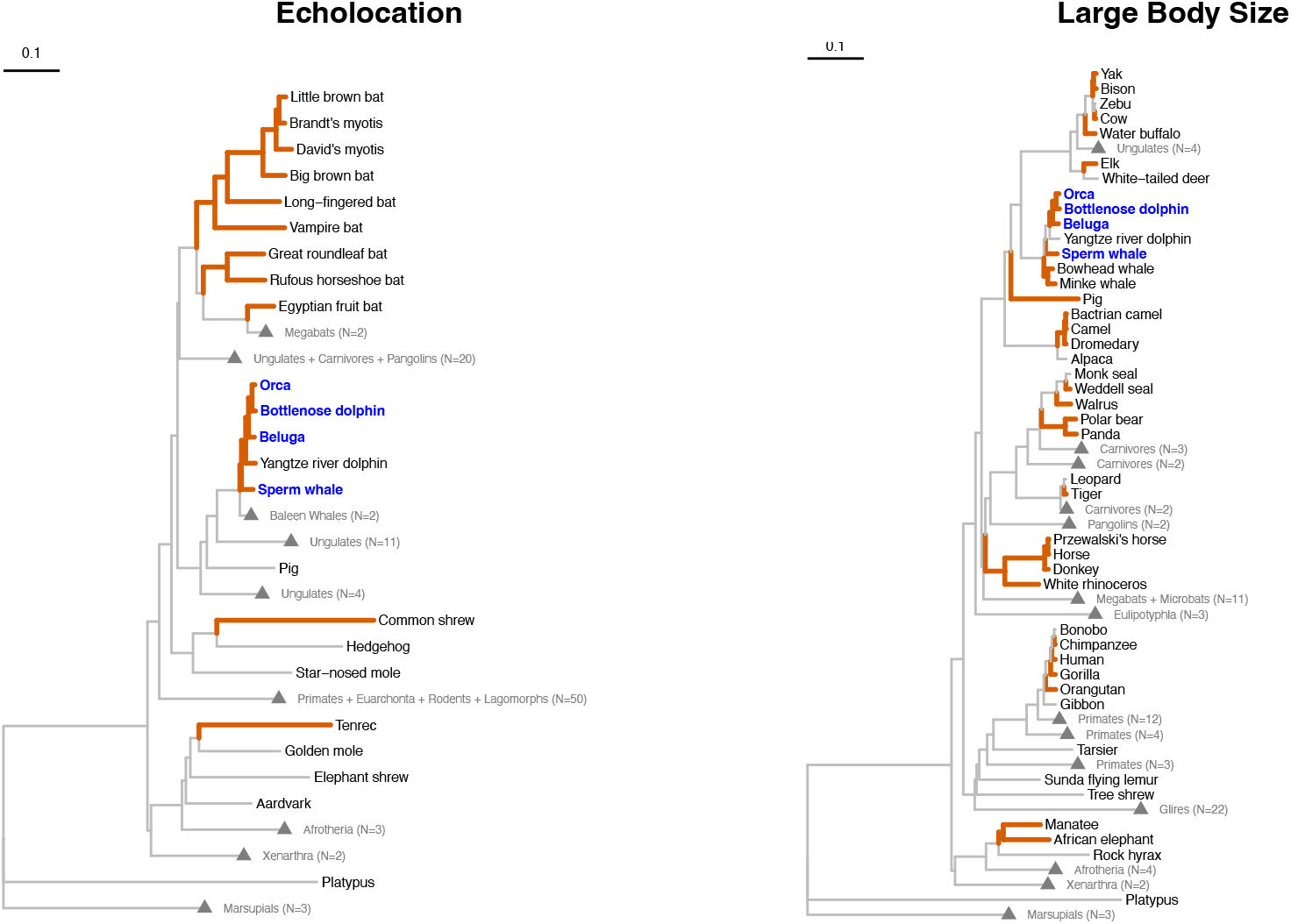
Phylogenetic trees for the echolocation (left) and large body size (right) studies. Phenotype-positive species and internal nodes where all descendants share the phenotype (foreground) are highlighted in orange. Species present in the foreground partitions of both analyses are highlighted in bold blue. Background clades lacking the phenotype are collapsed and labeled with their taxonomic group and species count.

`BUSTED-PH` employs a Union-Intersection test framework to evaluate a composite hypothesis. Biologically, we wish to identify genes where positive selection is (1) present in the foreground and (2) distinct from the background regime (Table 1). This corresponds to rejecting a composite null hypothesis that either there is no foreground selection, or that the foreground and background regimes are indistinguishable.

To assert a trait association, we must reject this composite null. Statistically, this is implemented as an Intersection-Union Test (IUT), which requires the simultaneous rejection of the component null hypotheses. We therefore compute a combined *p*-value, *p*_*c*_ = max (*p*_1_, *p*_3_), where *p*_1_ tests for foreground selection (Model 1 vs 2) and *p*_3_ tests for regime difference (Model 1 vs 4). This formulation ensures that *p*_*c*_ is a statistically valid (though conservative) *p*-value for the joint requirement of both conditions, preserving the nominal Type I error rate [Berger and Hsu, 1996].

In genome-wide scans, we apply standard multiple-testing corrections (e.g., Benjamini-Hochberg) directly to the vector of *p*_*c*_ values across all analyzed genes. This ensures that the primary discovery phase correctly accounts for the massive number of hypotheses tested. Furthermore, to distinguish specific trait-association from pervasive global selection, we examine the background lineages. A non-significant background selection test (*p*_2_) is informative but effectively “accepting the null,” which requires caution. We therefore apply an AIC-based model selection criterion. If the Akaike Information Criterion (AIC) prefers the null model (constrained background, Model 3) over the alternative (unconstrained, Model 1)—specifically if the LRT statistic for Test 2 is *<* 4—we infer that the background lacks strong evidence of selection.

We therefore define the protocol for identifying trait-associated positive selection as follows:

1. Compute *p*_*c*_ = max (*p*_1_, *p*_3_) for each gene.
2. Apply Benjamini-Hochberg FDR correction to the set of all *p*_*c*_ values.
3. Identify genes where corrected *p*_*c*_ ≤ *α* (e.g., 0.05).
4. For these significant hits, evaluate the background selection test specificity filter. The statistical significance of the trait association is determined solely by the Intersection-Union Test (*p*_*c*_ ≤ 0.05), which requires rejecting both the null of no foreground selection and the null of shared selective regimes. This ensures that the foreground distribution is statistically distinguishable from the background. We subsequently apply a heuristic filter based on the background selection signal (LRT for Test 2 *<* 4) to prioritize candidates where the selective signal is exclusive to the trait, rather than merely intensified relative to a background that is also under selection. This threshold of 4 corresponds to an AIC difference greater than 2 (since the alternative model has 2 additional parameters), indicating that the constrained model is statistically preferred (evidence ratio > 2.7) or indistinguishable from the unconstrained one.

### Functional enrichment and network modularity analysis

To characterize the biological functions and phenotypic consequences of the candidate gene sets identified by BUSTED-PH (Echolocation and Body Size), we performed a dual-pronged analysis integrating automated gene set enrichment and protein-protein interaction (PPI) network topology.

#### Automated Gene Set Enrichment (Enrichr)

We performed an automated over-representation analysis (ORA) using the Enrichr platform [Chen et al., 2013, Kuleshov et al., 2016]. This tool aggregates data from multiple annotated gene libraries, allowing us to cross-reference our candidate lists against diverse biological categories. We queried the following libraries: GO Biological Process 2023 to identify overrepresented biological pathways and molecular processes; MGI Mammalian Phenotype (Level 4, 2021) to link candidate genes to specific phenotypic abnormalities observed in mouse knockout models (e.g., “head tossing” as a proxy for vestibular dysfunction); Jensen TISSUES to determine if the candidate genes are preferentially expressed in specific tissues or cell types (e.g., “Outer hair cell”); and the GWAS Catalog 2019 to assess overlap with genes associated with human complex traits.

Enrichment significance was assessed using Fisher’s exact test. *P* -values were adjusted for multiple hypothesis testing using the Benjamini-Hochberg False Discovery Rate (FDR) method. Terms with an adjusted *P* -value *<* 0.05 were considered statistically significant.

#### Protein-Protein Interaction (PPI) Modularity

To test whether the candidate genes interact physically or co-functionally, we performed a permutation-based network modularity test. We utilized the human protein-protein interaction network from the STRING database (v12.0) [Szklarczyk et al., 2023]. Only interactions with a medium confidence score (≥ 400) were included to ensure robustness.

The connectivity of the candidate gene set was quantified by the number of observed edges (direct interactions) between mapped genes in the network (*E*_*obs*_). To determine statistical significance while controlling for network topology biases, we performed a degree-preserving permutation test. We generated a null distribution by randomly sampling 1,000 gene sets of the same size as the candidate list from the background universe. For each candidate gene, a random gene was selected from the background set that shared the same node degree (number of connections) in the STRING network. For each random set, we calculated the number of edges (*E*_*rand*_). The empirical *P* -value was calculated as (*r* + 1)*/*(*n* + 1), where *r* is the number of random sets with *E*_*rand*_ ≥ *E*_*obs*_, and *n* = 1000. A Z-score was also computed to quantify the effect size. This approach ensured that the observed connectivity was not an artifact of high-degree “hub” genes being overrepresented in our candidate lists.

### Empirical data analysis

#### Application to established positive controls

To validate the performance of `BUSTED-PH` on well-characterized instances of molecular adaptation, we analyzed three canonical positive control genes, previously demonstrated to exhibit convergent or lineage-specific adaptive evolution linked to phenotypic traits. For each gene, published multiple sequence alignments and associated binary (presence/absence) trait annotations were utilized. We applied `BUSTED-PH` to test for episodic diversifying selection (EDS) associated with the specified trait, with `BUSTED-E` employed to account for residual alignment error.

For *Prestin* (*SLC26A5* ) and the ATP alpha-1 subunit (*ATP1* ), alignments and foreground/background annotations were obtained from [Fukushima and Pollock, 2023]. For *SEMG2*, the alignment and phenotype mapping were sourced from the the `TraitRELAX` study [Halabi et al., 2021]. Species annotations are visualized in Figure S1. All analyses were executed using `HyPhy` version 2.5.94 or later, incorporating the flags `–error-sink Yes` and `–rates 2`.

#### Genome-wide analysis of echolocation in mammals

To identify genes potentially associated with the evolution of echolocation in mammals, we applied `BUSTED-PH` to a genome-wide dataset of orthologous protein-coding genes from 120 mammalian species, comprising both echolocating and non-echolocating lineages [Hecker and Hiller, 2020]. Species were annotated as echolocating (foreground) or non-echolocating (background), as depicted in Figure (left) [Gould, 1965, Holland et al., 2004, Liu et al., 2014]; internal branches were labeled using the **Conjunctive** strategy, which assigns the foreground state to an ancestral node only if all its descendants possess the trait. We generated protein-coding sequence alignments by extracting protein-coding regions from the 120-species whole-genome alignment based on UCSC ‘canonical’ human gene models for genome assembly hg38 [Hecker and Hiller, 2020]. Exons were extracted from the MAF alignment file using the `sub.msa` function of the `RPHAST` package [Hubisz et al., 2011], and the human reading frame was enforced via the `codon.clean.msa` function within the same package.

#### Genome-wide analysis of large body size in mammals

To further assess the utility of `BUSTED-PH` in detecting trait-associated selection, we conducted an additional analysis focusing on the evolution of large body size using the same 120-species mammalian whole-genome alignment source [Hecker and Hiller, 2020]. Species were annotated as “large” (foreground) if their reported body mass exceeded 50 kg, with all remaining species classified as the background (Figure 1 (right)). This threshold captures a diverse group of 37 large mammals, including cetaceans (e.g., *Physeter catodon, Orcinus orca*), pinnipeds (e.g., *Odobenus rosmarus, Leptonychotes weddellii* ), large ungulates (e.g., *Ceratotherium simum, Bos taurus, Camelus ferus*), carnivores (e.g., *Ursus maritimus, Panthera tigris*), and great apes (e.g., *Gorilla gorilla, Homo sapiens*), distinct from the echolocation trait. While the 50 kg cutoff is heuristic, it effectively isolates lineages that have undergone significant gigantism relative to the median mammalian body size. As with the echolocation analysis, we applied the **Conjunctive** strategy for ancestral branch labeling.

### Software availability

`BUSTED-PH` is implemented within the `HyPhy` software package (version 2.5.73 or later, we used version 2.5.94, as this is the first version implementing the Union-Intersection testing framework) [Kosakovsky Pond et al., 2020]. The analysis is executed using the `hyphy busted-ph` command. Interactive notebooks have been developed to facilitate the visualization of `BUSTED-PH` results (https://observablehq.com/@hyphy/busted-ph), including a dedicated resource for the genome-wide mammalian analyses (https://observablehq.com/@spond/busted-ph-analysis-summary). These notebooks provide a platform for users to upload and visualize their own `BUSTED-PH` JSON output files.

## Results

### Application to established positive controls

To evaluate the performance of `BUSTED-PH`, we applied the method to three canonical positive control genes with well-documented evidence of adaptive evolution associated with specific phenotypic traits. These genes served to verify whether `BUSTED-PH` could recover expected signals of episodic diversifying selection (EDS) linked to binary traits. We defined evidence of EDS by the following criteria: (i) detection of positive selection on foreground branches, (ii) absence of positive selection on background branches, and (iii) a statistically significant difference between the foreground and background selective regimes. We conducted analyses using two non-error *ω* rate classes (*K* = 2), as the *K* = 3 model resulted in overfitting (indicated by zero-weight or collapsed rate classes) confirmed by AIC_c_ comparisons.

#### Prestin (SLC26A5)

This gene is critical for high-frequency auditory sensitivity in echolocating bats [Li et al., 2008] and marine mammals [Li et al., 2010], representing a seminal example of molecular convergence. Applying `BUSTED-PH` to the alignment and phenotype annotations provided by [Fukushima and Pollock, 2023] (Table 2), we detected a clear signal of EDS in echolocating lineages. We inferred approximately 1% of the alignment to be subject to EDS on the foreground branches, with no evidence of selection on the background branches and a significant difference between the two *ω* distributions. Consistent with recent findings [Selberg et al., 2025], this alignment exhibited residual error on background lineages, as illustrated in Figure S2.

**Table 2.** Exemplar genes previously analyzed for evidence of convergent evolution. K = 2 rate classes were used for all analyses. Foreground* indicates the test for selection on foreground branches (Test 1). Background indicates the test for selection on background branches (Test 2). Shared indicates the test for difference in selective regimes (Test 3). ω_E_ denotes the error sink rate class (proportion).

| Gene | LRT $p$-value | $\omega_1$ (weight) | $\omega_2$ (weight) | $\omega_E$ (weight) | Branches (L) |
| --- | --- | --- | --- | --- | --- |
| ***Prestin*** (echolocation, 128 sequences, 758 codons) |  |  |  |  |  |
| Foreground* | **0.015** | 0.09 (98.8%) | 7.15 (1.2%) | N.D. | 19 (0.19) |
| Background | 0.50 | 0.04 (96.9%) | 1.06 (3.1%) | 201 (0.043%) | 228 (3.13) |
| Shared | < 0.001 | 0.05 (98.8%) | 2.60 (1.2%) | 287 (0.028%) | 247 (3.32) |
| ***ATP1A1*** (cardiac glycoside resistance, 29 sequences, 1045 codons) |  |  |  |  |  |
| Foreground* | **0.001** | 0.01 (99.3%) | 5.20 (0.7%) | N.D. | 20 (0.71) |
| Background | 0.50 | 0.01 (98.3%) | 1.00 (1.7%) | 203 (0.115%) | 35 (7.11) |
| Shared | < 0.001 | 0.01 (98.3%) | 1.00 (1.7%) | 100 (0.052%) | 55 (7.82) |
| ***SEMG2*** (polygynandry, 23 sequences, 586 codons) |  |  |  |  |  |
| Foreground* | < 0.001 | 1.00 (59.0%) | 4.58 (40.8%) | 100 (0.24%) | 8 (0.72) |
| Background | 0.20 | 1.00 (29.9%) | 1.24 (70.1%) | N.D. | 33 (0.34) |
| Shared | < 0.001 | 0.57 (42.9%) | 2.25 (57.1%) | N.D. | 41 (1.06) |

#### AŁP alpha-1 subunit (AŁP1A1)

Convergent substitutions in the alpha-1 subunit of the Na^+^/K^+^-ATPase are associated with resistance to cardiac glycosides, such as ouabain, across diverse insect taxa [Dobler et al., 2012, Mohammadi et al., 2022]. Analysis of the *ATP1A1* alignment from [Fukushima and Pollock, 2023 ], using `BUSTED-PH` revealed a strong signal of EDS restricted to resistant species. We inferred approximately 1% of the alignment to be subject to EDS on foreground branches, while background lineages showed pervasive conservation. We confirmed a marked difference in selective regimes between foreground and background branches. We also identified residual alignment errors on background lineages [Selberg et al., 2025] (Figure S2).

#### SEMG2

Accelerated evolution and diversifying selection have been frequently reported for *SEMG2*, a structural component of the semen coagulum, in species with high levels of sperm competition. We applied `BUSTED-PH` to *SEMG2* sequences annotated for polyandry [Halabi et al., 2021]. We observed a robust signal of EDS in polyandrous species, with a substantial fraction (approximately 40%) of the alignment subject to positive selection. Selective regimes differed significantly between foreground and background branches. We detected residual alignment error on the foreground lineages in this dataset [Selberg et al., 2025] (Figure S2).

### Simulations

We evaluated the performance of `BUSTED-PH` across twenty-one simulation scenarios, assessing Type I and Type II error rates and robustness to model misspecification.

These simulations evaluate `BUSTED-PH` performance when the evolutionary model is correctly specified.

#### Nectar-consuming birds

We obtained a phylogeny of 45 bird species from [Osipova et al., 2025], designating four nectar-consuming clades as the foreground (Figure S3). We generated three null (N) and ten power (P) simulation scenarios (Table 3; Birds scenarios). Key observations include:

**Table 3.**
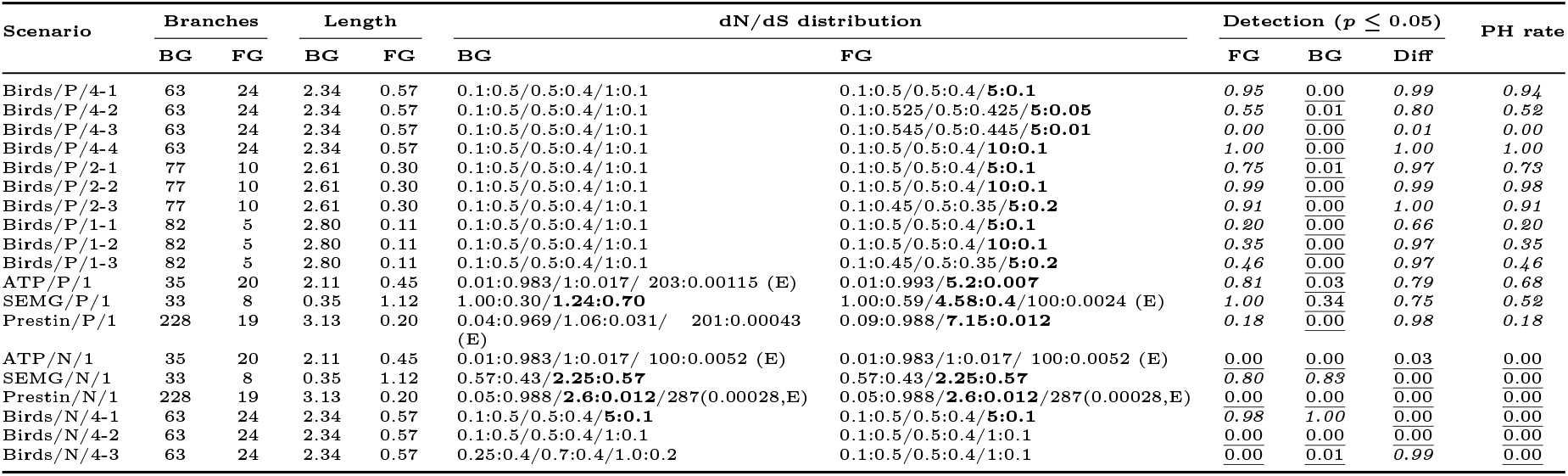
`BUSTED-PH` performance on data simulated under the correct model (BS-REL), using either the reference tree from Osipova et al. [Osipova et al., 2025] or Shultz and Sackton [Shultz and Sackton, 2019] with all (**/4**), two (**/2**) or one (**/1**) nectar consuming clades specified as foreground; or using one of the three datasets from Table 2 as templates for replicate generation. **/P** denote power simulations, **/N** denote null simulations. BG: background, FG: foreground. **Branches**: the number of tree branches in the corresponding set. **Length**: the cumulative length (subs/site) of branches in the corresponding set. **dN/dS distribution**: the distribution of dN/dS applied to the corresponding branch set; values with dN/dS > 1 are shown in bold. **Detection**: the rate of detection (/100 replicates) for the following tests, each at p ≤ 0.05 (following within-alignment Holm-Bonferroni correction); **FG**: EDS on foreground branches, **BG**: EDS on background branches, **Diff**: dN/dS distributions are different between FG and BG. Values in italics are expected to be high, values that are underlined are expected to be low. **PH rate**: the fraction of alignments where the BUSTED-PH composite test holds (EDS on FG, no EDS on BG, significant difference between FG and BG).

1. `BUSTED-PH` exhibited well-controlled false positive rates (N/4 simulations) both in the absence of EDS on foreground branches (N/4-2, N/4-3) and when EDS was present on both background and foreground branches (N/4-1).
2. Power to detect EDS on foreground lineages varied with:
  a. Effect size: Power was robust (PH rate > 0.90) when the proportion of sites under selection was sufficient (e.g., ≥ 10% at *ω* = 5), but declined significantly as the proportion dropped (e.g., P/4-2, P/4-3).
  b. Number of foreground clades: Detection improved with an increasing number of independent origins of the phenotype. Scenarios with four foreground clades (P/4) yielded higher PH rates (up to 1.00) compared to single-clade scenarios (P/1), which had PH rates ≤ 0.46.
3. The regime difference test proved critical for specificity. In scenarios with background selection (e.g., N/4-1), this test successfully differentiated regimes (Diff detection = 0.00), effectively filtering out potential false positives despite high detection rates by the foreground selection test alone (FG detection = 0.98).
4. `BUSTED-PH` proved robust to moderate background selection; in scenario N/4-1, *ω* > 1 on a subset of background branches did not induce spurious trait associations (PH rate = 0.00).

Overall, `BUSTED-PH` demonstrates high specificity and sufficient power when the number of independent phenotypic transitions and the strength of selection are adequate.

#### Exemplar genes

We utilized the topologies, branch lengths, substitution biases, codon frequencies, and site counts from the three exemplar genes in Table 2. We simulated 100 replicates using the exact inferred *ω* distributions (P/1 scenarios) for foreground and background branches, and 100 replicates employing a single shared *ω* distribution across the entire tree (Shared model, N/1 scenarios). In the latter case, detection rates for selective differences between branches were predictably low. In the former case, we observed the following:

#### 1. Prestin

Power was limited (18/100) due to the low simulated proportion of *ω >* 1 on foreground branches (∼1%) and the short total length of these branches (0.2 substitutions). However, the method demonstrated nearly perfect power (98/100) to detect regime differences. The overall `BUSTED-PH` detection rate was limited by the foreground power.

#### 2. ATP

Foreground selection detection was high (81/100). As expected, detection on background branches was low (3/100) given the absence of simulated EDS. The power to detect regime differences was high (79/100), resulting in a respectable overall detection rate of 68%.

#### 3. SEMG2

Foreground detection was robust (100/100), reflecting the substantial fraction (40%) of the alignment under positive selection. However, because weak positive selection was also simulated on the background (*ω* = 1.24), 34% of replicates reported selection on the background, reducing the overall power to 52%. This case is notable; the empirical result suggested weak background selection, a signal effectively recovered in the simulations.

#### Model misspecification

We analyzed data simulated with `evolver` [Yang, 2007], utilizing parameters from [Kowalczyk et al., 2021]. As Table 4 indicates, `BUSTED-PH` found no evidence of association with a trait under the pervasive selection scenario (PAML/N/1) but correctly identified association under the selective regime where only the foreground is under EDS (PAML/P/1). Thus, the method intrinsically accounts for non-specific selection, a scenario previously requiring “drop-out” testing. Furthermore, as the simulation models differ from the `BUSTED-PH` random effects model, these results demonstrate a degree of model robustness.

**Table 4.** BUSTED-PH performance on data simulated under a model that’s different from the BS-REL model, drawn from previous publications [ ]. **/P** denote power simulations, **/N** denote null simulations. BG: background, FG: foreground. **Branches**: the number of tree branches in the corresponding set. **Length**: the cumulative length (subs/site) of branches in the corresponding set. **dN/dS distribution**: the distribution of dN/dS applied to the corresponding branch set; values with dN/dS > 1 are shown in bold. **Detection**: the rate of detection (/100 replicates) for the following tests, each at p ≤ 0.05 (following within-alignment Holm-Bonferroni correction); **FG**: EDS on foreground branches, **BG**: EDS on background branches, **Diff**: dN/dS distributions are different between FG and BG. Values in italics are expected to be high, values that are underlined are expected to be low. **PH rate**: the fraction of alignments where the BUSTED-PH composite test holds (EDS on FG, no EDS on BG, significant difference between FG and BG).

| Scenario | Branches | | Length | | dN/dS distribution | | Detection ($p \leq 0.05$) | | | PH rate |
| --- | --- | --- | --- | --- | --- | --- | --- | --- | --- | --- |
|  | BG | FG | BG | FG | BG | FG | FG | BG | Diff |  |
| PAML/P/1 | 10 | 3 | 3.85 | 1.15 | 0.1:0.1/1.0:0.5 | 0.1:0.05/1.0:0.5/**5.0:0.9** | 1.00 | 0.00 | 1.00 | 1.00 |
| PAML/N/1 | 10 | 3 | 3.85 | 1.15 | 0.1:0.05/1.0:0.5/**5.0:0.9** | 0.1:0.05/1.0:0.5/**5.0:0.9** | 1.00 | 1.00 | 0.00 | 0.00 |

### Genome-wide analysis of echolocation in mammals

We analyzed a total of 19,153 orthologous gene alignments from a 120-species mammalian whole-genome alignment [Hecker and Hiller, 2020], with echolocation defined as the binary phenotype of interest. Echolocation represents a canonical system for studying molecular convergence, with seminal research established over two decades ago [Li et al., 2010, Parker et al., 2013, Liu et al., 2014]. More recently, echolocation-associated traits and genes have been utilized as benchmark datasets for the development of comparative genomic methods [Kowalczyk et al., 2019].

The dataset encompasses three distinct mammalian lineages, representing aquatic, airborne, and terrestrial forms of echolocation. The Cetaceans (*Physeter catodon, Tursiops truncatus*, and *Orcinus orca*) represent the sophisticated aquatic clade, utilizing specialized phonic lips to generate powerful nasal clicks. Within the Chiroptera, the dataset includes two distinct evolutionary strategies: sophisticated laryngeal echolocators, ranging from high-duty cycle species (*Rhinolophus*) to low-duty cycle vesper bats (*Myotis*). The Egyptian fruit bat (*Rousettus aegyptiacus*) serves as a unique evolutionary exception within the Pteropodidae, having independently developed lingual (tongue-click) echolocation. Finally, terrestrial insectivores from the Afrotheria (*Echinops telfairi* ) and Eulipotyphla (*Sorex araneus*) clades are included; these species possess a comparatively ancestral or rudimentary form of echolocation, utilizing simple tongue clicks or ultrasonic squeaks for close-range spatial orientation.

We annotated a total of 16 species as echolocating. To define the full foreground set, we employed the **Conjunctive** labeling strategy, where internal branches are included only if all their descendants are echolocators. This approach allows us to test for continued adaptive evolution throughout the history of the echolocating clades while remaining conservative compared to parsimony-based reconstruction. This resulted in the inclusion of up to 11 additional internal branches (Figure 1 (left)). Due to variation in species coverage across alignments, the number of testable branches differed between genes. A total of *N* = 18, 953 alignments (Figure 2) contained at least one echolocating species and were thus amenable to analysis. `BUSTED-PH` is not computationally light; it takes approximately 1 hour per gene on 4 ARM Neoverse cores, meaning that the entire genome-wide analysis finishes in a few days on a small lab cluster. However, this performance compares favorably with many existing methods: standard branch-site models in `PAML` are significantly slower, while Bayesian frameworks and recent convergence metrics such as *ω*_*C*_ [Fukushima and Pollock, 2023] also carry high computational demands. Consequently, `BUSTED-PH` represents a computationally reasonable solution for large-scale phylogenomic discovery.

**Figure 2.**
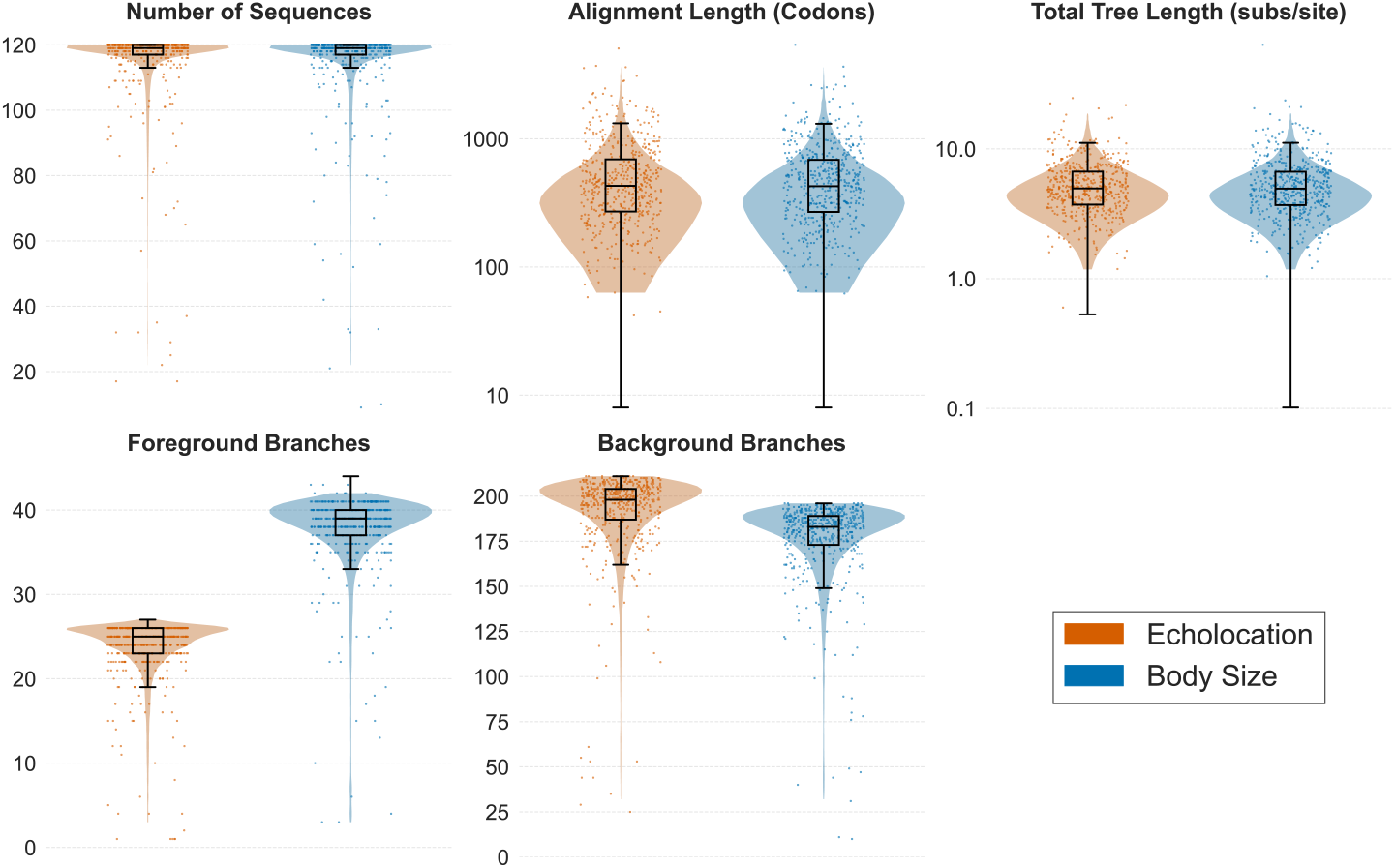
Descriptive statistics of the alignments analyzed for echolocation (N=18,953) and large body size (N=19,056). Distributions are shown for the number of species sequences, alignment length (codons), total tree length (substitutions/site), and the number of foreground (FG) and background (BG) branches. Note the logarithmic scale for alignment and tree lengths. For visual clarity, the density (violin) component of the plots excludes the extreme 0.5% tails of each distribution.

Applying a False Discovery Rate (FDR) of 5% (Benjamini-Hochberg) and requiring evidence of neutral or constrained evolution on background lineages, we identified 72 genes where EDS was significantly associated with the echolocating trait (Table S2, Figure 3). As illustrated in Figure 3, the `BUSTED-PH` protocol acts as a rigorous filter: while 761 genes show evidence of selection on the echolocating foreground branches alone, the majority are excluded because they either lack a significant difference from the background regime or exhibit pervasive selection across the phylogeny. This winnowing process is critical for isolating trait-specific signals from broader genomic background noise. To characterize the biological cohesion of these 72 candidates, we performed protein-protein interaction (PPI) modularity analysis and automated gene set enrichment (Enrichr). PPI analysis revealed that the candidate set does not form a statistically significant interaction module (*P* = 0.06), suggesting that the trait evolved via adaptive changes in multiple distinct systems rather than a single tight protein complex. Consistent with this, functional enrichment identified specific, disparate biological signals. Significant enrichment was observed in the MGI Mammalian Phenotype ontology for “head tossing” (*P* = 0.038), a behavioral proxy for vestibular and inner ear dysfunction in mice, and in the Jensen TISSUES library for “Outer hair cell” (*P* = 0.037), directly linking the candidate set to the sensory machinery of echolocation.

**Figure 3.**
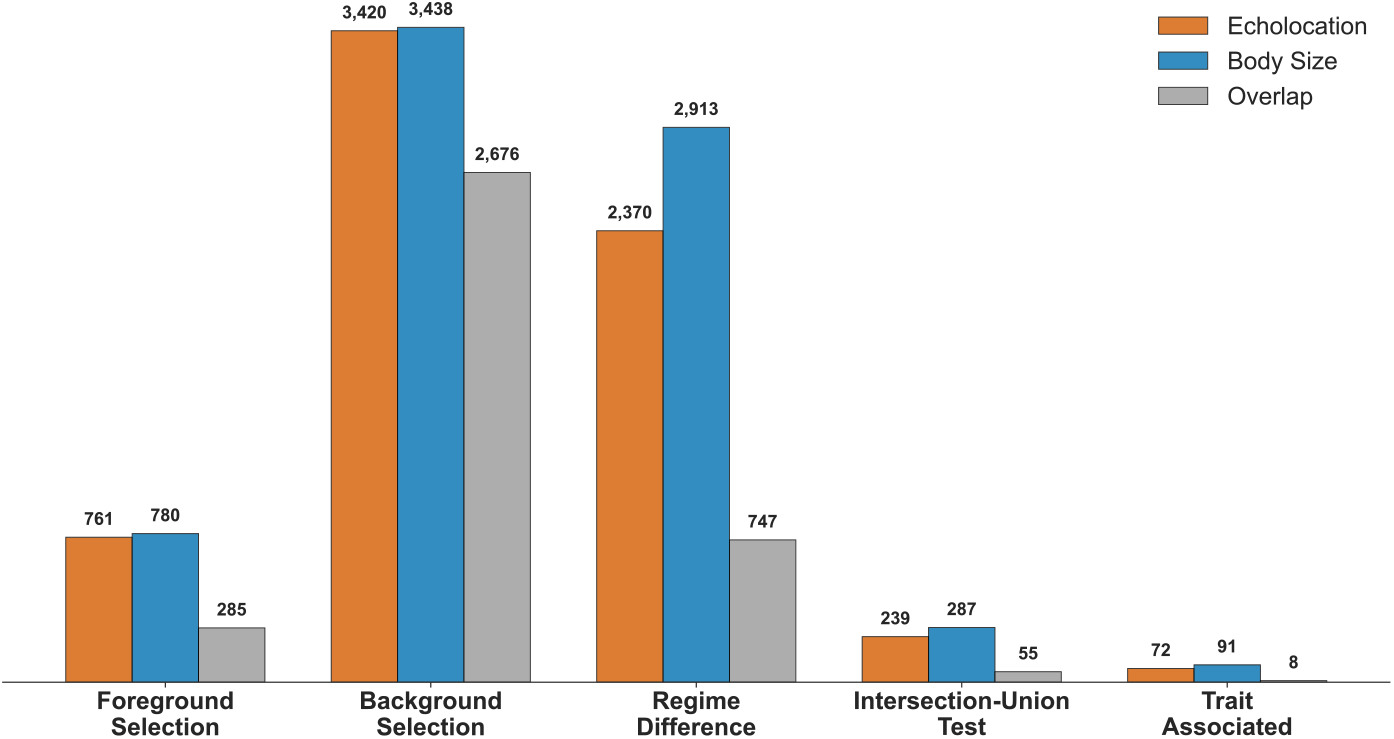
Comparison of selection results for Echolocation (orange) and Large Body Size (blue). The bar chart displays the number of genes identified by each component of the `BUSTED-PH` test: evidence of episodic diversifying selection (EDS) on foreground branches (Test 1), background branches (Test 2), significant difference between foreground and background selective regimes (Test 3), the intersection of foreground selection and regime difference (Intersection-Union Test), and the final set of Trait Associated genes (Intersection-Union Test filtered for lack of strong background selection). The grey bars indicate the number of genes shared between the two traits for each category.

While these functional enrichments align with broad physiological expectations, we note that standard enrichment analyses on small gene sets derived from comparative genomic scans generally warrant cautious interpretation. Such analyses are frequently biased towards well-characterized pathways (e.g., auditory function in humans and mice) and may fail to capture novel or lineage-specific adaptations where gene ontology annotations are sparse or generic [Stoeger et al., 2018]. Furthermore, the statistical power of enrichment tests is inherently limited when the input list is small (*N* = 72), potentially masking subtler functional modules that contribute to the complex polygenic phenotype of echolocation [Khatri et al., 2012]. The detection of 72 genes associated with echolocation serves as both a validation of the `BUSTED-PH` pipeline and a source of novel evolutionary insights. We organize these candidates into three broad categories: core sensory and neural machinery, physiological support systems, and likely ecological confounders driven by the distinct life histories of echolocating lineages.

#### Core Sensory and Neural Machinery

The strongest signal comes from genes governing auditory perception and neural circuit formation. We recover canonical echolocation drivers *SLC26A5* (Prestin), *TMC1*, and *CDH23*, alongside *KARS1, TMEM63C*, and the newly identified *CHD7* and *SCN4B* (sodium channel beta subunit). While *CHD7* is linked to CHARGE syndrome and ear abnormalities in humans, its pleiotropic role in chromatin remodeling suggests it may contribute to the regulation of otic development in echolocators, though the precise mechanism remains to be determined. Beyond the ear, the expanded list reveals a robust signature of neural adaptation. Selection on *GRIN2A* (NMDA receptor), *SOX3* (neural progenitor marker), *MYT1L* (neuronal differentiation), and *EPHB2* (axon guidance) points to the specialized neural development required to process complex acoustic spatial information.

#### Physiological Adaptations

Echolocation imposes unique metabolic and physiological demands. We detect selection on lipid homeostasis and fatty acid sensing genes (*ABCA13, HSD17B13, FFAR3, GPR42* ), likely facilitating the maintenance of acoustic fats in cetacean melons and jaws. Additionally, *ACE* (blood pressure regulation) may reflect adaptations to the extreme hemodynamic stresses of diving or flight [Xu et al., 2013].

#### Ecological and Life-History Confounders

Several candidates likely reflect the environmental constraints of aquatic (cetacean) or aerial (bat) niches rather than echolocation itself. Genes involved in skin and barrier integrity (*KRT83, LCE1C, LCE2D, ADAMTS7, ECM1, SPRR5* ) and immunity (*CGAS, TAX1BP1, SIGLECL1, TERF2IP, CLEC7A, CXCL5, CXCL6* ) appear to track with the integumentary and metabolic challenges of these lifestyles (“Flight-as-Fever”) [Marcovitz et al., 2019, Langlois, 2025, O’Shea et al., 2014]. Similarly, a cluster of testis-enriched genes (*ACTBL2, MAGED4B, SPATA31D4, KLHL10, ADGRG2* ) likely emerges from the generally elevated evolutionary rates driven by sexual selection in these lineages [Swanson and Vacquier, 2002].

#### Comparison with previously reported echolocating genes

A substantial body of literature has proposed a suite of “key echolocating genes” based on varying statistical criteria. Even with a relaxed FDR of 5%, which expands our candidate list to 72 genes, `BUSTED-PH` consistently recovers core drivers like *SLC26A5, TMC1*, and *CDH23* while omitting others such as *OTOF* and *PCDH15*. This persistence of discordance is instructive; it confirms that the discrepancy is not merely a matter of statistical thresholding but reflects the fundamental difference in the hypothesis being tested. We require evidence of episodic diversifying selection that is *specific* to the echolocating foreground and distinct from the background.

Many genes previously identified using standard branch-site models (e.g., in `PAML`) may capture pervasive selection or relaxation rather than trait-specific adaptation. For instance, while early studies reported convergence in *PCDH15* and *OTOF* [Shen et al., 2012], subsequent analyses controlling for neutral homoplasy challenged these findings [Parker et al., 2013, Lambert et al., 2017]. Parallel evolution in *KCNQ4* has also been supported by site-specific methods [Liu et al.,2011, Chai et al., 2020]. Methods relying on tree incongruence, such as the Site-Specific Log-Likelihood Support (SSLS) used by [Parker et al.,2013], identify convergence by comparing the fit of alignments to the species tree versus alternative convergent topologies. In several cases, these convergent substitutions are sufficiently numerous to drive gene-tree topologies that conflict with the established species phylogeny, grouping echolocators together [Liu et al., 2011, Chai et al., 2020]. However, *KCNQ4* remains absent from our results; this suggests that while specific codons undergo convergent substitution, the gene-wide selective regime does not shift significantly enough to meet the `BUSTED-PH` criteria. By explicitly contrasting foreground and background regimes, `BUSTED-PH` likely filters out these ambiguous or localized signals. Conversely, our recovery of *SLC26A5, TMC1*, and *CDH23* aligns with robust consensus across methods, including recent sparse learning approaches [Allard et al., 2025], confirming these as core adaptive loci where selective regimes shift definitively with the phenotype.

### Genome-wide analysis of large body size in mammals

Analysis of a total of *N* = 19, 056 alignments (Figure 2) for large body size (>50 kg) using the **Conjunctive** labeling strategy identified 91 genes with significant evidence of trait-associated episodic diversifying selection Table S3. This candidate list is comparable in scale to the echolocation-associated gene set (72 genes) and reflects molecular adaptations to the physical and physiological constraints of gigantism.

#### Musculoskeletal and Structural Reinforcement

Large body size imposes increased mechanical stress, scaling with the square-cube law. Identified genes include extracellular matrix components (*LAMA2, LAMA5, HMCN2* ), the synovial lubricant *PRG4* [Jay et al., 2007], and nuclear envelope proteins (*SYNE2, TOR1AIP2* ) that maintain nuclear structural integrity under high-torque muscle contractions [Zhang et al.,2001]

#### Regulation of Organ and Skeletal Growth

Adaptations in developmental scaling are evidenced by selection on *WWC3*, a regulator of the Hippo signaling pathway that limits organ size [Genevet et al., 2010, Chen et al., 2020]. Selection was also identified in *CUL7, SMC1A*, and *TUBGCP6*. As mutations in these genes are associated with growth disorders in humans (e.g., primordial dwarfism), their divergence in large mammals may reflect modifications to fundamental growth regulatory pathways [Huber et al., 2005, Deardorff et al., 2007, Puffenberger et al., 2012].

#### Physiological Scaling and Cardiac Function

Allometric scaling of metabolic and cardiovascular systems is reflected in selection on metabolic capacity genes (*GCLC, SOAT1, FADS3* ) and cardiac regulators. Selection on *SCN4B* [Medeiros-Domingo et al., 2007] and *TMEM94* [Stephen et al., 2018] likely relates to the specialized cardiac dynamics required to support large body volumes and stabilized heart rates.

#### Genomic Integrity and Cancer Suppression

Consistent with the theoretical requirement to suppress cancer in large-bodied, long-lived species (Peto’s Paradox [Abegglen et al., 2015]), we identified selection on several genomic safeguards. These include *PLK3*, involved in cell-cycle arrest and DNA damage detection [Xie et al., 2001], *TERF2IP*, which protects chromosome ends [Sfeir et al., 2010], and the chromatin remodeler *RUVBL1* [Jha et al., 2008].

#### Enrichment and Overlap

In contrast to the echolocation results, PPI analysis indicated that the 91 body size candidates form a significantly interacting functional module (*P* = 0.004). This suggests that despite the broad systemic challenges of gigantism, the underlying molecular adaptations are integrated into a cohesive protein interaction network. Automated enrichment analysis identified “Sensory Perception of Smell” (GO:0007608, *P* = 0.042) as the only significant category. This result likely reflects ecological confounding rather than adaptation for size: many large-bodied mammals in our foreground (e.g., cetaceans) have lost or reduced olfaction, leading to relaxed selection on olfactory receptors which `BUSTED-PH` detects as a regime shift relative to terrestrial background species. Indeed, approximately 16% of the list comprises olfactory receptors (*OR* families) and zinc-finger proteins, which we interpret as a specific signature of the aquatic transition in the cetacean foreground. Finally, eight genes (*SCN4B, RUVBL1, EPC2, TAX1BP1, KARS1, TMEM63C, TERF2IP, SRRM1* ) were common to both the echolocation and body size analyses, possibly reflecting the shared presence of cetaceans in both foregrounds, or the influence of other shared traits (Figure 4).

**Figure 4.**
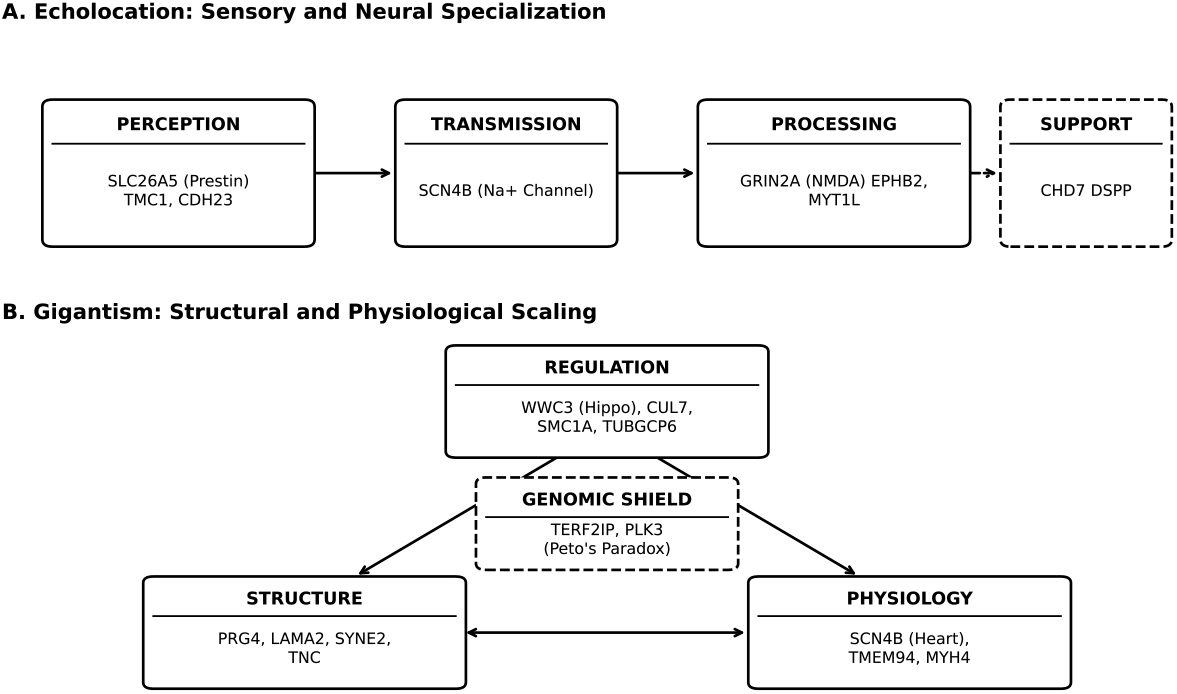
Overview of key functional modules identified by `BUSTED-PH`. (A) Genes associated with echolocation map to the entire sensory signal chain, from auditory perception and neural processing to morphological support. (B) Genes associated with mammalian gigantism cluster into regulatory, structural, and physiological roles, with a central “shield” of genomic integrity genes protecting against cancer (Peto’s Paradox). Solid boxes represent direct functional drivers of the phenotype, while dashed boxes denote foundational developmental or permissive support mechanisms.

### Data and code availability

All data and code utilized in this study are publicly available. The multiple sequence alignments analyzed are hosted at https://github.com/nclark-lab/ComparativeData/wiki. `BUSTED-PH` is distributed as part of the `HyPhy` package (version 2.5.94), which may be downloaded from, along with supplementary processing scripts. Technical documentation for the analyses is provided at https://github.com/veg/hyphy-readmes. Interactive notebooks for data visualization are accessible at https://observablehq.co Accessibility will be maintained to ensure the reproducibility of all findings.

## Discussion

We introduce `BUSTED-PH`, a branch-site codon model designed to identify genes where episodic diversifying selection (EDS) is associated with a binary trait. By explicitly modeling and contrasting positive selection across phenotype-defined foreground (FG) and background (BG) branches, `BUSTED-PH` advances beyond standard branch-site methods. Our analyses of positive controls, genome-wide scans, and extensive simulations demonstrate that `BUSTED-PH` offers a statistically rigorous and biologically interpretable framework for detecting trait-associated adaptation—a capability that is increasingly vital as projects like Zoonomia and the Vertebrate Genomes Project (VGP) populate the tree of life with high-quality genomic data.

Application to three canonical genes—*Prestin, ATP1A1*, and *SEMG2* —validated the model’s capacity to recover established instances of convergent adaptation. In all cases, `BUSTED-PH` detected significant EDS restricted to foreground lineages, characterized by a significant difference between foreground and background selective regimes. These results underscore the sensitivity of the method and its robustness to residual alignment errors, accounted for by the integrated `BUSTED-E` component [Selberg et al., 2025]. This integration is particularly crucial for large-scale comparative genomics, where automated pipelines may leave residual misalignment or annotation errors that would otherwise generate false positives.

Our genome-wide scan for echolocation-linked adaptation identified 72 genes with significant evidence of trait-associated EDS. These candidates cluster into functional modules associated with auditory perception and neural processing. Selection on *SLC26A5* (Prestin), *TMC1*, and *CDH23* is consistent with molecular adaptations of the cochlear amplifier and hair bundle mechanics for high-frequency reception. Convergence in *ACTBL2* (stereocilia cytoskeleton) provides further evidence of hair bundle modification. Potential adaptations for temporal processing include *SCN4B*, associated with rapid channel gating, and *GRIN2A*, involved in synaptic plasticity. Neural development candidates *EPHB2* and *MYT1L* suggest involvement of axonal guidance pathways. Morphological changes are implied by *CHD7* (otic capsule development) and *DSPP*, which is linked to mandibular bone density in toothed whales. Physiological candidates like *CGAS* and *RUVBL1* may relate to metabolic or stress responses associated with this lifestyle.

Extending our framework to the evolution of large body size, we identified 91 genes exhibiting convergent positive selection in large-bodied lineages (Table S3). Similar to the echolocation analysis, `BUSTED-PH` filtered a large initial set of genes (780 with foreground selection) down to 91 high-confidence candidates (Figure 3). This stringent reduction (approximately 88%) underscores the importance of controlling for background selection and pervasive regime shifts, ensuring that the identified genes reflect specific adaptations to large body size rather than generalized evolutionary trends. While “large body size” is a complex, composite trait influenced by diverse ecological factors (e.g., aquatic buoyancy vs. terrestrial gravity), our screen effectively targets the correlates of gigantism that transcend these specific niches. The functional profiles of these genes suggest adaptations to the universal physical and physiological constraints of increased mass. Selection on *CUL7* and *WWC3* implies modifications to developmental pathways regulating organ size and skeletal growth. Biomechanical constraints are reflected in the selection of structural and extracellular matrix components, including *PRG4, LAMA2*, and *SYNE2*, potentially mitigating increased physical stress. Furthermore, the identification of tumor suppressors (*PLK3, TERF2IP* ) and metabolic regulators (*SCN4B, MYH4* ) aligns with the physiological requirements of large body size, specifically relating to cancer suppression (Peto’s Paradox) and metabolic scaling. These results indicate that the evolution of gigantism involves specific molecular changes in structural, developmental, and physiological systems. While phenotype-associated shifts in selective pressure can manifest as relaxation of purifying selection rather than positive selection, `BUSTED-PH` explicitly tests for sites with *ω >* 1, thereby distinguishing adaptive diversifying selection from relaxation. Other methods in the `HyPhy` package, such as `RELAX` [Wertheim et al., 2015], are specifically designed to test for the latter.

Simulation studies reinforced these empirical findings. `BUSTED-PH` maintained strict false positive control across null scenarios, even in the presence of background selection. Power to detect foreground-specific EDS correlated with the number of independent phenotypic origins and the strength of selection. Crucially, the test for a difference between foreground and background *ω* distributions emerged as the definitive filter for specificity, effectively distinguishing trait-associated signals from pervasive selection.

`BUSTED-PH` occupies a distinct niche in the methodological landscape (Table S4). While rate-based methods like `RERconverge` [Kowalczyk et al., 2019] excel at detecting gene-wide shifts—often signaling relaxed selection—they may lack the resolution for site-specific adaptive bursts. Conversely, methods seeking identical amino acid substitutions, including the error-corrected *ω*_*C*_ metric [Fukushima and Pollock, 2023], face theoretical limitations imposed by the “Stokes Shift” [Pollock et al.,2012 ] and can be confounded by functional equivalence. Emerging approaches using protein language models, such as ACEP [ Cao et al., 2025, ],offer a promising avenue for detecting high-order convergence but currently retain the computational overhead of traditional models for significance testing. By focusing on the *process* of evolution (shifts in selective pressure) rather than the *outcome*, `BUSTED-PH` overcomes these constraints, detecting genes where the *regime* of selection converges, even if the specific mutations differ. Furthermore, our use of the Conjunctive labeling strategy explicitly tests for the maintenance or early origin of the trait in ancestral lineages, providing a conservative test for deep homology, though it may miss convergent events restricted solely to the tips. This methodological distinction underscores a broader philosophical trade-off in comparative genomics. Methods like `BUSTED-PH` prioritize the identification of a high-confidence set of “top” candidates through rigorous hypothesis testing and strict false discovery rate control, making them ideal for pinpointing specific molecular drivers. Conversely, many study designs prioritize power over specificity, seeking to rank thousands of genes to uncover aggregate pathway-level signals, often tolerating a higher false positive rate at the individual gene level. While `BUSTED-PH` produces rankings suitable for enrichment analyses, its primary strength lies in its capacity to filter pervasive background noise, thereby isolating specific trait-associated adaptation.

Despite its advantages, `BUSTED-PH` is not without limitations. First, as a gene-wide test for episodic selection, it primarily detects signals where multiple sites or strong selective bursts occur; consequently, it may lack the power to identify genes driven by only a few directional substitutions. Second, like all comparative methods that assume a fixed species tree, it may be susceptible to false positives arising from gene tree discordance (hemiplasy) [Mendes and Hahn, 2016]. However, `BUSTED-PH` can be applied equally well to individual gene trees, and we encourage researchers to consider this approach. While running analyses on both species and gene topologies is twice as computationally expensive, the contrast can be highly informative for disentangling true convergent adaptation from ILS-driven artifacts. Third, the method is subject to phenotypic confounding. If a set of lineages designated with the trait of interest (e.g., echolocation) also shares a correlated but unmodeled trait (e.g., specialized skin adaptations or flight), `BUSTED-PH` may identify selection that is actually driven by the confounder. However, our multi-stage testing procedure progressively mitigates this effect: for instance, while 285 genes show significant foreground selection in both the echolocation and large body size analyses, this overlap is reduced to 55 genes following the Intersection-Union Test, and finally to just eight high-confidence genes after applying the background specificity filter (Figure 3). This shared signal likely reflects the prominent role of cetaceans in both foregrounds, driving signals in genes like *SCN4B* and *RUVBL1* that are biologically plausible for both traits, yet the rapid decline in overlap demonstrates the protocol’s effectiveness at winnowing out broader lineage-specific signals in favor of trait-associated adaptation. Fourth, the choice of the background branch set involves a critical trade-off. Utilizing a background that is too small or closely related may underpower the detection of pervasive selection, while employing large alignments with highly divergent species increases the likelihood of alignment errors, which can manifest as false positive signals despite error-sink mitigation. Finally, the current implementation relies on binary phenotype annotations, potentially oversimplifying complex, continuous, or multi-state traits that might be better modeled by frameworks capable of handling quantitative data. Sixth, we note that like all codon models, `BUSTED-PH` may be susceptible to confounding by GC-biased gene conversion (gBGC), particularly in mammals, although the requirement for foreground-specific shifts helps mitigate this if gBGC is uniform across the tree.

By providing a unified framework that explicitly tests for selection in both foreground and background lineages and statistically contrasts their regimes, `BUSTED-PH` enhances the precision of comparative genomic studies seeking to unravel the genetic basis of complex convergent traits.

## References

L. M. Abegglen, A. F. Caulin, A. Chan, et al. Potential mechanisms for cancer resistance in elephants and comparative cellular response to dna damage in humans. JAMA, 314:1850–1860, 2015.

J. B. Allard, S. Sharma, R. Patel, M. Sanderford, K. Tamura, S. Vucetic, G. S. Gerhard, and S. Kumar. Evolutionary sparse learning reveals the shared genetic basis of convergent traits. Nat. Commun., 16:3217, 2025. doi: 10.1038/s41467-025-58428-8.

D. W. Armitage, A.G. Alonso-Sánchez, S. R. Coy, Z. Cheng, A. Hagenbeek, K.P. López-Martínez, Y. H. Phua, and A. R. Sears. Adaptive pangenomic remodeling in the azolla cyanobiont amid a transient microbiome. ISME J., 19:wraf154, 2025.

M. Barkdull and C. S. Moreau. Worker reproduction and caste polymorphism impact genome evolution and social genes across the ants. Genome Biol. Evol., 15:evad095, 2023.

C. A. Berger, D. K. Steinberg, L. A. Copeman, and A. M. Tarrant. Comparative analysis of the molecular starvation response of southern ocean copepods. Mol. Ecol., 34:e17371, 2025.

R. L. Berger and J. C. Hsu. Bioequivalence trials, intersection-union tests and equivalence confidence sets. Stat. Sci., 11:283–319, 1996.

Z. Cao, H. Zhang, and Z. Zou. Language models reveal a complex sequence basis for adaptive convergent evolution of protein functions. Proc. Natl. Acad. Sci., 122:e2418254122, 2025.

S. Chai, R. Tian, X. Rong, G. Li, B. Chen, W. Ren, S. Xu, and G. Yang. Evidence of echolocation in the common shrew from molecular convergence with other echolocating mammals. Zool. Stud., 59:e4, 2020. doi: 10.6620/ZS.2020.59-4.

Edward Y Chen, Christopher M Tan, Yan Kou, Qiaonan Duan, Zichen Wang, Gustavo V Meirelles, Neil R Clark, and Avi Ma’ayan. Enrichr: interactive and collaborative html5 gene list enrichment analysis tool. BMC Bioinformatics, 14(1):128, 2013.

Y. Chen, H. Han, G. Seo, R. E. Vargas, B. Yang, K. Chuc, H. Zhao, and W. Wang. Systematic analysis of the hippo pathway organization and oncogenic alteration in evolution. Sci. Rep., 10:3173, 2020.

P. A. Christin, N. Salamin, V. Savolainen, and G. Besnard. C4 photosynthesis evolved in grasses via parallel adaptive genetic changes. Curr. Biol., 17:1241–1247, 2007.

F. Cicconardi, C. F. McLellan, A. Seguret, W. O. McMillan, and S. H. Montgomery. Convergent molecular evolution associated with repeated transitions to gregarious larval behavior in heliconiini. Mol. Biol. Evol., 42:msaf179, 2025.

M. A. Deardorff, M. Kaur, D. Yaeger, et al. Mutations in cohesin complex members smc3 and smc1a cause a mild variant of cornelia de lange syndrome with predominant mental retardation. Am. J. Hum. Genet., 80:485–494, 2007.

R. R. Dhakal, A. Harkess, and P. G. Wolf. Chromosome numbers and reproductive life cycles in green plants: A phylotranscriptomic perspective. Plant Direct, 9:e70044, 2025.

S. Dobler, S. Dalla, V. Wagschal, and A. A. Agrawal. Community-wide convergent evolution in insect adaptation to toxic cardenolides by substitutions in the na,k-atpase. Proc. Natl. Acad. Sci., 109:13040– 13045, 2012.

M. L. Donaldson, M. Barkdull, and C. S. Moreau. Comparative genomics analyses reveal selection on neuronal and cuticular hydrocarbon genes is associated with aggression in ants (hymenoptera: Formicidae). Ann. Entomol. Soc. Am., 118:37–58, 2025.

S. Dong, X. Li, Q. Liu, T. Zhu, A. Tian, N. Chen, X. Tu, and L. Ban. Comparative genomics uncovers evolutionary drivers of locust migratory adaptation. BMC Genomics, 26:203, 2025.

K. Fukushima and D. D. Pollock. Detecting macroevolutionary genotype–phenotype associations using error-corrected rates of protein convergence. Nat. Ecol. Evol., 7:155–170, 2023.

A. Genevet, M. C. Wehr, R. Brain, et al. Kibra is a regulator of the salvador/warts/hippo signaling network. Dev. Cell, 18:300–308, 2010.

E. Gould. Evidence for echolocation in the tenrecidae of madagascar. Proc. Am. Philos. Soc., 109:352–360, 1965.

K. Halabi, E. L. Karin, L. Guéguen, and I. Mayrose. A codon model for associating phenotypic traits with altered selective patterns of sequence evolution. Syst. Biol., 70:608–622, 2021.

N. Hecker and M. Hiller. A genome alignment of 120 mammals highlights ultraconserved element variability and placenta-associated enhancers. GigaScience, 9:giz159, 2020.

R. A. Holland, D. A. Waters, and J. M. V. Rayner. Echolocation signal structure in the megachiropteran bat rousettus aegyptiacus geoffroy 1810. J. Exp. Biol., 207:4361–4369, 2004.

Z. Hu, T. B. Sackton, S. V. Edwards, and J. S. Liu. Bayesian detection of convergent rate changes of conserved noncoding elements on phylogenetic trees. Mol. Biol. Evol., 36:1086–1100, 2019.

C. Huber, D. Dias-Santagata, A. Glaser, J. O’Sullivan, R. Brauner, K. Wu, X. Xu, K. Pearce, R. Wang, M. L. G. Uzielli, et al. Identification of mutations in cul7 in 3-m syndrome. Nat. Genet., 37:1119–1124, 2005. doi: 10.1038/ng1628.

M. J. Hubisz, K. S. Pollard, and A. Siepel. Phast and rphast: phylogenetic analysis with space/time models. Brief. Bioinform., 12:41–51, 2011.

G. D. Jay, J. R. Torres, M. L. Warman, M. C. Laderer, and K. S. Breuer. The role of lubricin in the mechanical behavior of synovial fluid. Proc. Natl. Acad. Sci., 104:6194–6199, 2007.

S. Jha, E. Shibata, and A. Dutta. Human rvb1/tip49 is required for the histone acetyltransferase activity of tip60/nua4 and for the downregulation of phosphorylation on h2ax after dna damage. Mol. Cell. Biol., 28:2690–2700, 2008.

Purvesh Khatri, Marina Sirota, and Atul J Butte. Ten years of pathway analysis: current approaches and outstanding challenges. PLoS computational biology, 8(2):e1002375, 2012.

E. E. K. Kopania, G. W. C. Thomas, C. R. Hutter, S. M. E. Mortimer, C. M. Callahan, E. Roycroft, A. S. Achmadi, W. G. Breed, N. L. Clark, J. A. Esselstyn, et al. Sperm competition intensity shapes divergence in both sperm morphology and reproductive genes across murine rodents. Evolution, 79:11–27, 2025.

S. L. Kosakovsky Pond and S. D. W. Frost. Not so different after all: A comparison of methods for detecting amino acid sites under selection. Mol. Biol. Evol., 22:1208–1222, 2005.

S. L. Kosakovsky Pond, A. F. Y. Poon, R. Velazquez, S. Weaver, N. L. Hepler, B. Murrell, S. D. Shank, B. R. Magalis, D. Bouvier, A. Nekrutenko, et al. Hyphy 2.5—a customizable platform for evolutionary hypothesis testing using phylogenies. Mol. Biol. Evol., 37:295–299, 2020.

A. Kowalczyk, W. K. Meyer, R. Partha, W. Mao, N. L. Clark, and M. Chikina. Rerconverge: an r package for associating evolutionary rates with convergent traits. Bioinformatics, 35:4815–4817, 2019.

A. Kowalczyk, R. Partha, N. L. Clark, and M. Chikina. Pan-mammalian analysis of molecular constraints underlying extended lifespan. eLife, 9:e51089, 2020.

Amanda Kowalczyk, Maria Chikina, and Nathan L Clark. A cautionary tale on proper use of branch-site models to detect convergent positive selection. bioRxiv, 2021. doi: 10.1101/2021.10.26.465984. URL

Maxim V Kuleshov, Matthew R Jones, Andrew D Rouillard, Nicolas F Fernandez, Qiaonan Duan, Zichen Wang, Semyon Koplev, Scott L Jenkins, Katherine M Jagodnik, Alexander Lachmann, et al. Enrichr: a comprehensive gene set enrichment analysis web server 2016 update. Nucleic Acids Research, 44(W1): W90–W97, 2016.

M. J. Lambert, A. A. Nevue, and C. V. Portfors. Contrasting patterns of adaptive sequence convergence among echolocating mammals. Genome Biol. Evol., 9:44–57, 2017.

Ryan A. Langlois. Does fever drive the evolution of antiviral genes? Journal of Experimental Medicine, 223 (1):e20251747, 12 2025.

G. Li, J. Wang, S. J. Rossiter, G. Jones, J. A. Cotton, and S. Zhang. The hearing gene prestin reunites echolocating bats. Proc. Natl. Acad. Sci., 105:13959–13964, 2008.

Y. Li, Z. Liu, P. Shi, and J. Zhang. The hearing gene prestin unites echolocating bats and whales. Curr. Biol., 20:R55–R56, 2010.

Z. Liu, S. Li, W. Wang, D. Xu, R. W. Murphy, and P. Shi. Parallel evolution of kcnq4 in echolocating bats. PLOS ONE, 6:e26618, 2011.

Z. Liu, F.-Y. Qi, X. Zhou, H.-Q. Ren, and P. Shi. Parallel sites implicate functional convergence of the hearing gene prestin among echolocating mammals. Mol. Biol. Evol., 31:2415–2424, 2014.

Alexander G Lucaci, Jordan D Zehr, David Enard, Joseph W Thornton, and Sergei L Kosakovsky Pond. Evolutionary shortcuts via multinucleotide substitutions and their impact on natural selection analyses. Molecular Biology and Evolution, 40(7):msad150, 07 2023. ISSN 1537-1719. doi: 10.1093/molbev/msad150. URL https://doi.org/10.1093/molbev/msad150.

A. J. Ludington, J. M. Hammond, J. Breen, I. W. Deveson, and K. L. Sanders. New chromosome-scale genomes provide insights into marine adaptations of sea snakes (hydrophis: Elapidae). BMC Biol., 21:284, 2023.

A. Marcovitz, Y. Turakhia, H. I. Chen, M. Gloudemans, B. A. Braun, H. Wang, and G. Bejerano. A functional enrichment test for molecular convergent evolution finds a clear protein-coding signal in echolocating bats and whales. Nat. Ecol. Evol., 3:1803–1813, 2019.

A. Medeiros-Domingo, T. Kaku, D. J. Tester, et al. Scn4b-encoded sodium channel beta4 subunit in congenital long-qt syndrome. Circulation, 116:134–142, 2007.

F. K. Mendes and M. W. Hahn. Gene tree discordance can generate patterns of diminishing convergence over time. Mol. Biol. Evol., 33:3299–3307, 2016.

S. Mohammadi, S. Herrera-Álvarez, L. Yang, M. del P. Rodríguez-Ordoñez, K. Zhang, J. F. Storz, S. Dobler, J. Crawford, and P. Andolfatto. Constraints on the evolution of toxin-resistant na,k-atpases have limited dependence on sequence divergence. PLOS Genet., 18:e1010323, 2022.

Ariadna E Morales, Frank T Burbrink, Marion Segall, Maria Meza, Chetan Munegowda, Paul W Webala, Bruce D Patterson, Vu Dinh Thong, Manuel Ruedi, Michael Hiller, and Nancy B Simmons. Distinct genes with similar functions underlie convergent evolution in myotis bat ecomorphs. Mol. Biol. Evol., 41(9):msae165, 08 2024.

P. O. Mulhair, L. Crowley, D. H. Boyes, O. T. Lewis, and P. W. H. Holland. Opsin gene duplication in lepidoptera: Retrotransposition, sex linkage, and gene expression. Mol. Biol. Evol., 40:msad241, 2023.

B. Murrell, S. Weaver, M. D. Smith, J. O. Wertheim, S. Murrell, A. Aylward, K. Eren, T. Pollner, D. P. Martin, D. M. Smith, et al. Gene-wide identification of episodic selection. Mol. Biol. Evol., 32:1365–1371, 2015.

S. V. Muse and B. S. Gaut. A likelihood approach for comparing synonymous and nonsynonymous nucleotide substitution rates, with application to the chloroplast genome. Mol. Biol. Evol., 11:715–724, 1994.

C. A. Onetto, C. M. Ward, C. Varela, L. Hale, S. A. Schmidt, and A. R. Borneman. Genetic and phenotypic diversity of wine-associated hanseniaspora species. FEMS Yeast Res., 25:foaf031, 2025.

T. J. O’Shea, P. M. Cryan, A. A. Cunningham, A. R. Fooks, D. T. S. Hayman, A. D. Luis, A. J. Peel, R. K. Plowright, and J. L. N. Wood. Bat flight and zoonotic viruses. Emerg. Infect. Dis., 20:741–745, 2014.

Ekaterina Osipova, Meng-Ching Ko, Konstantin M. Petricek, Simon Yung Wa Sin, Thomas Brown, Sylke Winkler, Martin Pippel, Julia Jarrells, Susanne Weiche, Mai-Britt Mosbech, Fanny Taborsak-Lines, Chuan Wang, Orlando Contreras-Lopez, Remi-Andre Olsen, Philip Ewels, Daniel Mendez-Aranda, Andrea Gaede, Keren Sadanandan, Gabriel Weijie Low, Amanda Monte, Ninon Ballerstaedt, Nicolas M. Adreani, Lucia Mentesana, Auguste von Bayern, Alejandro Rico-Guevara, Scott V. Edwards, Carolina Frankl-Vilches, Heiner Kuhl, Antje Bakker, Manfred Gahr, Douglas L. Altshuler, William A. Buttemer, Michael Schupp, Maude W. Baldwin, Michael Hiller, and Timothy B. Sackton. Convergent and lineage-specific genomic changes shape adaptations in sugar-consuming birds. bioRxiv, 2025. doi: 10.1101/2024.08.30.610474. URL

J. Parker, G. Tsagkogeorga, J. A. Cotton, Y. Liu, P. Provero, E. Stupka, and S. J. Rossiter. Genome-wide signatures of convergent evolution in echolocating mammals. Nature, 502:228–231, 2013.

R. Partha, B. K. Chauhan, Z. Ferreira, J. D. Robinson, K. Lathrop, K. K. Nischal, M. Chikina, and N. L. Clark. Subterranean mammals show convergent regression in ocular genes and enhancers, along with adaptation to tunneling. eLife, 6:e25884, 2017.

D. D. Pollock, G. Thiltgen, and R. A. Goldstein. Amino acid coevolution induces an evolutionary stokes shift. Proc. Natl. Acad. Sci., 109:E1352–E1359, 2012.

S. K. Pond, W. Delport, S. V. Muse, and K. Scheffler. Correcting the bias of empirical frequency parameter estimators in codon models. PLOS ONE, 5:e11230, 2010.

X. Prudent, G. Parra, P. Schwede, J. G. Roscito, and M. Hiller. Controlling for phylogenetic relatedness and evolutionary rates improves the discovery of associations between species’ phenotypic and genomic differences. Mol. Biol. Evol., 33:2135–2150, 2016.

E. G. Puffenberger, R. N. Jinks, C. Sougnez, K. Cibulskis, R. A. Willert, N. P. Achilly, et al. Genetic mapping and exome sequencing identify variants associated with five novel diseases. PLoS One, 7:e28936, 2012. doi: 10.1371/journal.pone.0028936.

A. Rhie, S. A. McCarthy, O. Fedrigo, J. Damas, G. Formenti, S. Koren, M. Uliano-Silva, W. Chow, A. Fungtammasan, J. Kim, et al. Towards complete and error-free genome assemblies of all vertebrate species. Nature, 592:737–746, 2021.

I. L. Ruesink-Bueno, A. Drews, E. A. O’Connor, and H. Westerdahl. Expansion of mhc-iib has constrained the evolution of mhc-iia in passerines. Genome Biol. Evol., 16:evae236, 2024.

T. B. Sackton, P. Grayson, A. Cloutier, Z. Hu, J. S. Liu, N. E. Wheeler, P. P. Gardner, J. A. Clarke, A. J. Baker, M. Clamp, et al. Convergent regulatory evolution and loss of flight in paleognathous birds. Science, 364:74–78, 2019.

R. F. Sage. The evolution of c4 photosynthesis. New Phytol., 161:341–370, 2004.

A. Selberg, N. L. Clark, T. B. Sackton, S. V. Muse, A. G. Lucaci, S. Weaver, A. Nekrutenko, M. Chikina, and S. L. K. Pond. Minus the error: Testing for positive selection in the presence of residual alignment errors. eLife, 2025. Available from: https://elifesciences.org/reviewed-preprints/106921.

S. G. Self and K.-Y. Liang. Asymptotic properties of maximum likelihood estimators and likelihood ratio tests under nonstandard conditions. J. Am. Stat. Assoc., 82:605–610, 1987.

A. Sfeir, S. Kabir, M. van Overbeek, et al. Loss of rap1 induces telomere recombination in the absence of nhej or a dna damage signal. Science, 327:1657–1661, 2010.

Y.-Y. Shen, L. Liang, G.-S. Li, R. W. Murphy, and Y.-P. Zhang. Parallel evolution of auditory genes for echolocation in bats and toothed whales. PLOS Genet., 8:e1002788, 2012.

A. J. Shultz and T. B. Sackton. Immune genes are hotspots of shared positive selection across birds and mammals. eLife, 8:e41815, 2019.

A. Singh, N. S. Pope, and M.M. López-Uribe. Shifts in bee diet breadths are associated with gene gains and losses and positive selection across olfactory receptors. G3 GenesGenomesGenetics, 15:jkaf105, 2025.

J. Stephen, S. Maddirevula, S. Nampoothiri, J. D. Burke, M. Herzog, A. Shukla, et al. Bi-allelic tmem94 truncating variants are associated with neurodevelopmental delay, congenital heart defects, and distinct facial dysmorphism. Am. J. Hum. Genet., 103:948–967, 2018. doi: 10.1016/j.ajhg.2018.11.001.

Thomas Stoeger, Martin Gerlach, Richard I Morimoto, and Luís A Nunes Amaral. Large-scale investigation of the reasons why potentially important genes are ignored. PLoS biology, 16(9):e2006643, 2018.

J. F. Storz. Causes of molecular convergence and parallelism in protein evolution. Nat. Rev. Genet., 17: 239–250, 2016.

W. J. Swanson and V. D. Vacquier. Rapid evolution of reproductive proteins. Nat. Rev. Genet., 3:137–144, 2002.

Damian Szklarczyk, Rebecca Kirsch, Mikaela Koutrouli, Katerina Nastou, Farrokh Mehryary, Radja Hachilif, Annika L Gable, Tao Fang, Nadezhda T Doncheva, Sampo Pyysalo, Peer Bork, Lars J Jensen, and Christian von Mering. The STRING database in 2023: protein-protein association networks and functional enrichment analyses for any sequenced genome of interest. Nucleic Acids Res., 51(D1):sD638–D646, 2023. doi: 10.1093/nar/gkac1000.

J. O. Wertheim, B. Murrell, M. D. Smith, S. L. Kosakovsky Pond, and K. Scheffler. Relax: Detecting relaxed selection in a phylogenetic framework. Mol. Biol. Evol., 32:820–832, 2015.

S. R. Wisotsky, S. L. Kosakovsky Pond, S. D. Shank, and S. V. Muse. Synonymous site-to-site substitution rate variation dramatically inflates false positive rates of selection analyses: Ignore at your own peril. Mol. Biol. Evol., 37:2430–2439, 2020.

S. Xie, H. Wu, Q. Wang, et al. Plk3 functionally links dna damage to cell cycle arrest and apoptosis at least in part via the p53 pathway. J. Biol. Chem., 276:43305–43312, 2001.

S. Xu, Y. Yang, X. Zhou, J. Xu, K. Zhou, and G. Yang. Adaptive evolution of the osmoregulation-related genes in cetaceans during secondary aquatic adaptation. BMC Evol. Biol., 13:189, 2013.

Z. Yang. Paml 4: Phylogenetic analysis by maximum likelihood. Mol. Biol. Evol., 24:1586–1591, 2007.

L. H. Yusuf, Y. Saldívar Lemus, P. Thorpe, C. Macías Garcia, and M. G. Ritchie. Genomic signatures associated with transitions to viviparity in cyprinodontiformes. Mol. Biol. Evol., 40:msad208, 2023.

Q. Zhang, J. M. Skeie, K. Burridge, et al. Nesprins: a novel family of spectrin-repeat-containing proteins that characterize the nuclear membrane in multiple tissues. J. Cell Sci., 114:4485–4498, 2001.

Zoonomia Consortium. A comparative genomics multitool for scientific discovery and conservation. Nature, 587:240–245, 2020.

