## Supplemental Figure 1 for "BUSTED-PH: Isolating the genomic signatures of convergent phenotypes"

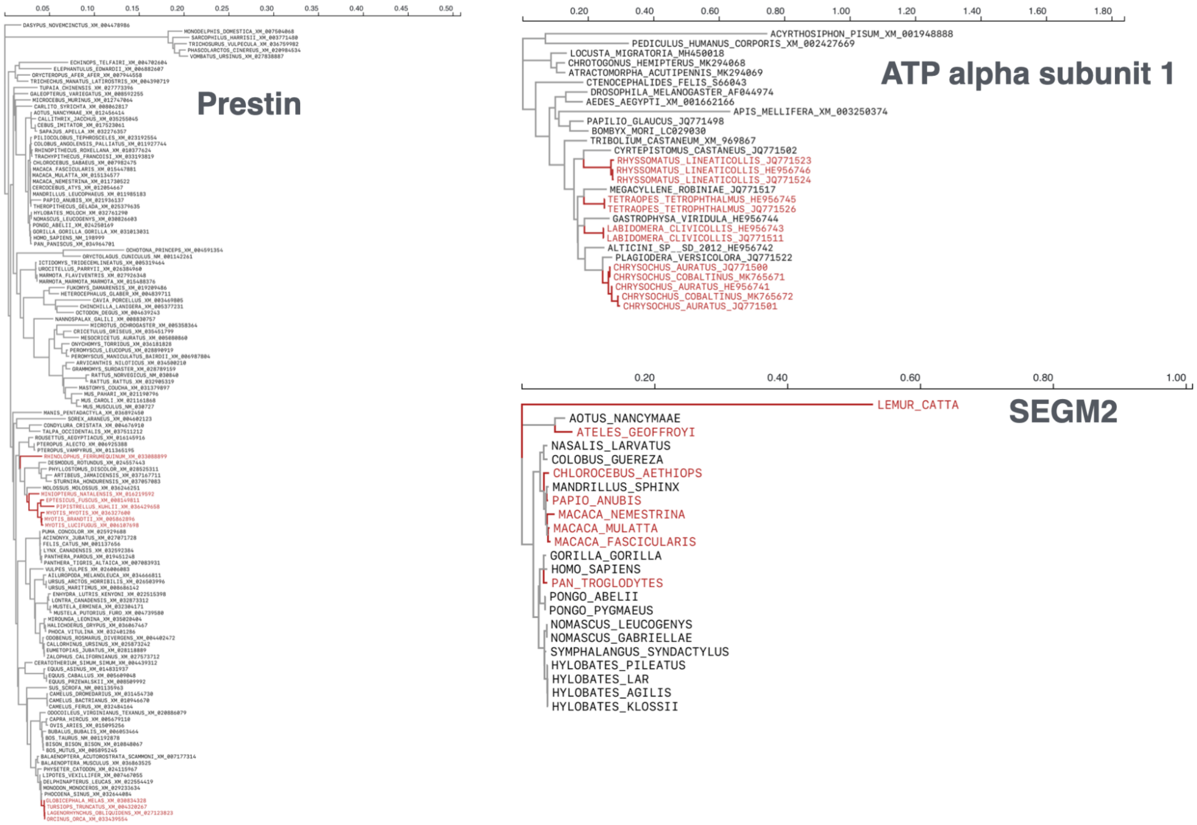


***Figure S1:*** *Phylogenetic trees with foreground branches shown in red for the three empirical datasets in Table 2.*
