## Supplemental Figure 2 for "BUSTED-PH: Isolating the genomic signatures of convergent phenotypes"

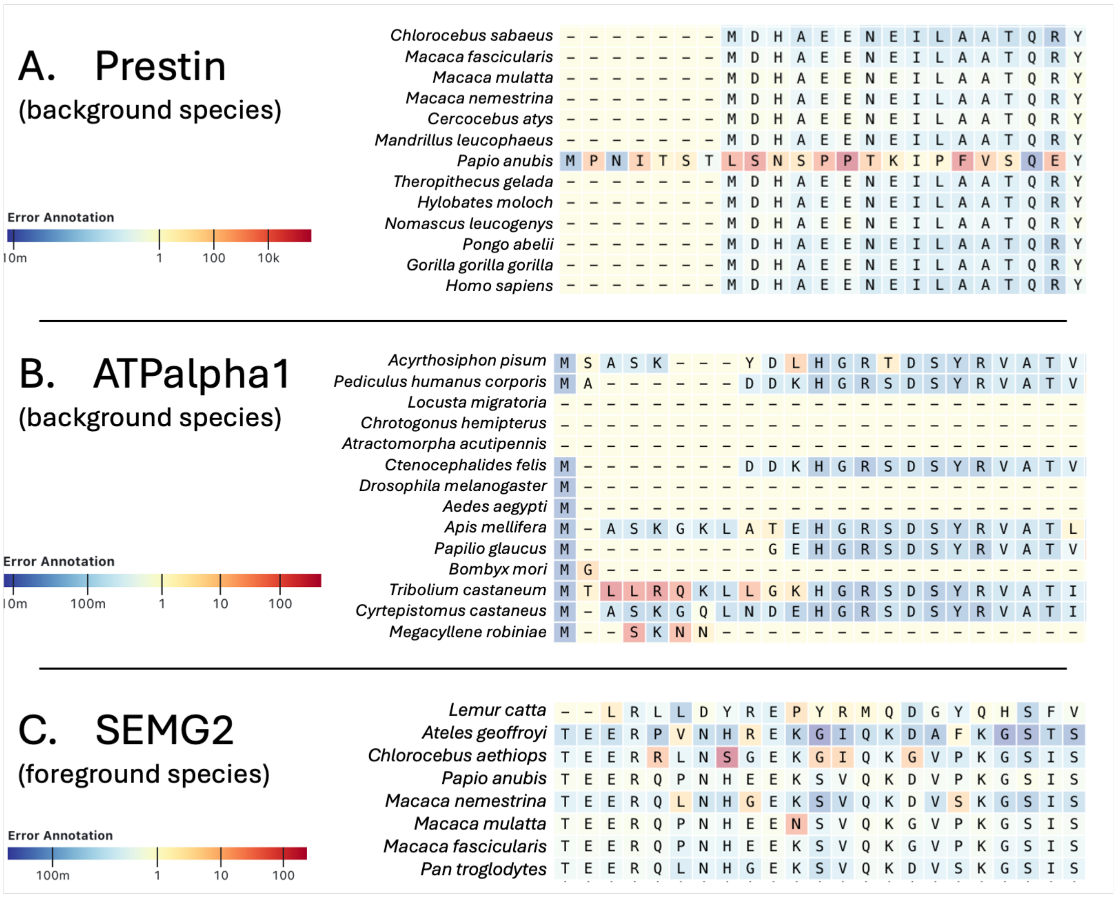


***Figure S2:*** *Residual alignment errors detected by* `BUSTED-PH` *in three positive-control genes. Prestin and ATPalpha1 (A, B) contain localized misaligned regions on background lineages, flagged by elevated error annotation values. SEMG2 (C) shows more scattered individual foreground codons inferred as highly unlikely substitutions. These examples reflect the residual errors present in the empirical alignments used for* `BUSTED-PH` *analyses.*
