## Supplementary figures and images for "BUSTED-PH: Isolating the genomic signatures of convergent phenotypes"

### Supplemental Figure 3

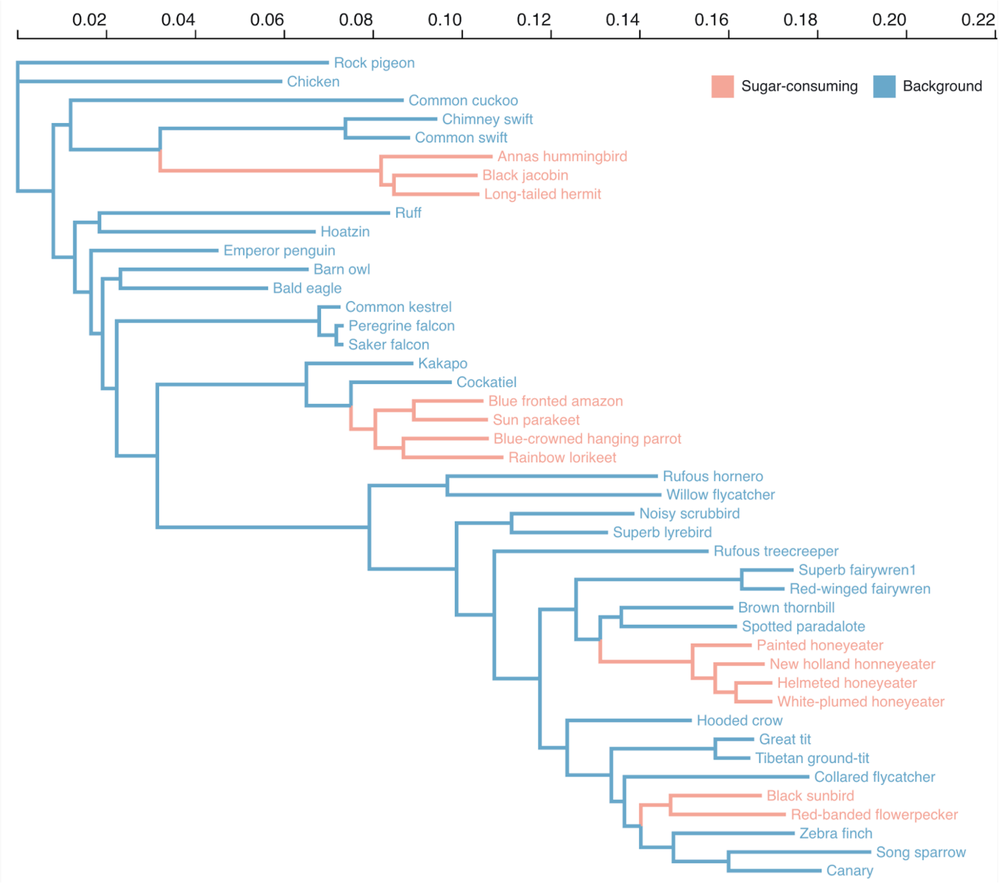


***Figure S3:*** *Tree used for power simulations (Table 3), from Osipova et al. [2025].*
