## Supplemental Table 1 for "BUSTED-PH: Isolating the genomic signatures of convergent phenotypes"

| Study | Organisms | Trait | Genes |
|---|---|---|---|
| Ludington et al. [2023] | Snakes | Marine adaptation | 8654 |
| Dong et al. [2025] | Locusts | Migratory behavior | 6874 |
| Cicconardi et al. [2025] | Heliconiini butterflies | Social behavior | 3393 |
| Dhakal et al. [2025] | Green plants | Chromosome number | 54 families |
| Ruesink-Bueno et al. [2024] | Birds | Passerine vs nonpasserine birds | 1; DAA1 |
| Onetto et al. [2025] | Yeast | Wine fermentation | 2036 |
| Armitage et al. [2025] | Symbiotic cyanobacteria | Azolla symbionts vs free-living bacteria | 3520 |
| Barkdull and Moreau [2023] | Ants | Worker caste polymorphism and reproductive capacity | 32792 |
| Singh et al. [2025] | Bees | Diet breadth | 251 |
| Berger et al. [2025] | Copepods | Oil sac used to sequester lipids | 13640 |
| Yusuf et al. [2023] | Ray-finned fishes | Viviparity vs oviparity | 1044 |
| Mulhair et al. [2023] | Lepidoptera | Nocturnal vs diurnal eye genes | 3 |
| Kopania et al. [2025] | Murine rodents | Relative testes mass | 11775 |
| Donaldson et al. [2025] | Ants | Aggression and nestmate discrimination | 16233 |
| bottomrule | | | |

***Table S1:*** *Studies that have implemented BUSTED-PH to test for lineage or trait-associated positive selection (excluding self-citations).*
