## Supplemental Table 2 for "BUSTED-PH: Isolating the genomic signatures of convergent phenotypes"

| Symbol | Gene | Function | Expression |
|---|---|---|---|
| ABCA13 | ATP binding cassette subfamily A member 13 | Lipid: Transport | Bone marrow |
| ACE | Angiotensin I converting enzyme | Metabolism: Blood pressure | Intestine, Testis |
| ACTBL2 | Actin beta like 2 | Sperm: Cytoskeleton | Not detected (HPA) |
| ADAMTS6 | ADAM metallopeptidase with thrombospondin type 1 motif 6 | ECM: Matrix degradation | Placenta |
| ADAMTS7 | ADAM metallopeptidase with thrombospondin type 1 motif 7 | ECM: Matrix degradation | Heart muscle, Blood vessel |
| ADGRG2 | Adhesion G protein-coupled receptor G2 | Sperm: Adhesion GPCR | Epididymis |
| C6ORF15 | Chromosome 6 open reading frame 15 | Skin: Follicle dev (likely) | Skin 1, Cervix, Lymphoid tis[illegible] |
| CDH23 | Cadherin related 23 | Auditory: Hair cell tip links | Inner Ear, Ovary |
| CGAS | Cyclic GMP-AMP synthase | Immunity: DNA sensor | Bone marrow |
| CHD7 | Chromodomain helicase DNA binding protein 7 | Auditory/Neural: CHARGE syndrome | Brain |
| CLEC7A | C-type lectin domain containing 7A | Immunity: Fungal recognition | Bone marrow, Lymphoid tiss[illegible] |
| CRYBG1 | Crystallin beta-gamma domain containing 1 | Cytoskeleton: Beta-gamma crystallin | Placenta, Esophagus |
| CXCL5 | C-X-C motif chemokine ligand 5 | Immunity: Neutrophil chemoattractant | Salivary gland, Lymphoid tis[illegible] |
| CXCL6 | C-X-C motif chemokine ligand 6 | Immunity: Neutrophil chemoattractant | Lymphoid tissue, Gallbladde[illegible] |
| DIS3L | DIS3 like exosome 3'-5' exoribonuclease | RNA: Exosome complex | Ubiquitous |
| DSPP | Dentin sialophosphoprotein | Teeth: Dentin mineralization | Teeth (Odontoblasts) |
| ECM1 | Extracellular matrix protein 1 | Skin: Extracellular matrix | Esophagus, Epididymis |
| EFCAB8 | EF-hand calcium binding domain 8 | Sperm: Calcium binding | Testis |
| EPC2 | Enhancer of polycomb homolog 2 | Regulation: Chromatin remodeling | Ubiquitous |
| EPHB2 | EPH receptor B2 | Neural: Axon guidance | Intestine |
| F11 | Coagulation factor XI | Blood: Coagulation | Liver |
| FBXO43 | F-box protein 43 | Cell cycle: Meiosis arrest | Testis |
| FBXO6 | F-box protein 6 | Ubiquitin: Glycoprotein quality | Ubiquitous |
| FFAR3 | Free fatty acid receptor 3 | Metabolism: Fatty acid receptor | Adipose tissue, Lymphoid tis[illegible] |
| FOXJ2 | Forkhead box J2 | Regulation: Transcription factor | Ubiquitous |
| FREM2 | FRAS1 related extracellular matrix 2 | Dev: Epithelial remodeling | Kidney, Thyroid gland |
| GOLGA2 | Golgin A2 | Transport: Golgi matrix | Ubiquitous |
| GPR161 | G protein-coupled receptor 161 | Signaling: GPCR (Cilia) | Smooth muscle, Endometriu[illegible] |
| GPR42 | G protein-coupled receptor 42 | Metabolism: Fatty acid receptor | Not detected (HPA) |
| GRIN2A | Glutamate ionotropic receptor NMDA type subunit 2A | Neural: NMDA receptor | Brain |
| HEATR4 | HEAT repeat containing 4 | Unknown: Heat repeat | Testis |
| HNRNPLL | Heterogeneous nuclear ribonucleoprotein L like | Splicing: Regulation | Ubiquitous |
| HSD17B13 | Hydroxysteroid 17-beta dehydrogenase 13 | Lipid: Droplet metabolism | Liver |
| ITIH1 | Inter-alpha-trypsin inhibitor heavy chain 1 | Inflammation: Trypsin inhibitor | Liver |
| KARS1 | Lysyl-tRNA synthetase 1 | Auditory: Protein synthesis (Deafness) | Ubiquitous |
| KLHL10 | Kelch like family member 10 | Sperm: Ubiquitination | Testis |
| KRT83 | Keratin 83 | Skin: Hair keratin | Skin 1 |
| LCE1C | Late cornified envelope 1C | Skin: Barrier formation | Skin 1 |
| LCE2D | Late cornified envelope 2D | Skin: Barrier formation | Skin 1 |
| MAGED4B | MAGE family member D4B | Sperm: Antigen | Brain, Pituitary gland |
| MYT1L | Myelin transcription factor 1 like | Neural: Neuronal differentiation | Brain, Pituitary gland |
| NDST4 | N-deacetylase and N-sulfotransferase 4 | Metabolism: Heparan sulfate | Not detected (HPA) |
| NUP62CL | Nucleoporin 62 C-terminal like | Transport: Nuclear pore | Epididymis, Fallopian tube |
| OR10T2 | Olfactory receptor family 10 subfamily T member 2 | Smell: Olfactory receptor | Not detected (HPA) |
| OR2F2 | Olfactory receptor family 2 subfamily F member 2 | Smell: Olfactory receptor | Not detected (HPA) |
| OR4C11 | Olfactory receptor family 4 subfamily C member 11 | Smell: Olfactory receptor | Not detected (HPA) |
| PADI2 | Peptidyl arginine deiminase 2 | Regulation: Citrullination | Tongue, Skeletal muscle, Bra[illegible] |
| PCDHGB1 | Protocadherin gamma subfamily B, 1 | Neural: Circuit formation | Brain |
| PCDHGB5 | Protocadherin gamma subfamily B, 5 | Neural: Protocadherin | Parathyroid gland |

| Symbol | Gene | Function | Expression |
|---|---|---|---|
| PPTC7 | Protein phosphatase targeting COQ7 | Metabolism: Phosphatase | Tongue |
| RUVBL1 | RuvB like AAA ATPase 1 | Regulation: Chromatin remodeling | Ubiquitous |
| SCN4B | Sodium voltage-gated channel beta subunit 4 | Auditory/Neural: Sodium channel beta | Tongue, Skeletal muscle, Bra[illegible] |
| SF3B1 | Splicing factor 3b subunit 1 | Splicing: Spliceosome | Ubiquitous |
| SIGLECL1 | SIGLEC family like 1 | Immunity: Sialic-acid binding | Testis |
| SLC26A5 | Solute carrier family 26 member 5 | Auditory: Outer hair cell motor | Inner Ear (Outer hair cells) |
| SNRNP200 | Small nuclear ribonucleoprotein U5 subunit 200 | Splicing: Helicase | Ubiquitous |
| SOX3 | SRY-box transcription factor 3 | Neural: Progenitor marker | Fallopian tube, Brain, Testis |
| SPATA31D4 | SPATA31 subfamily D member 4 | Sperm: Spermatogenesis | Testis |
| SPRR5 | Small proline rich protein 5 | Skin: Cornification | Skin 1 |
| SRRM1 | Serine and arginine repetitive matrix 1 | Splicing: RNA processing | Ubiquitous |
| STON1-GTF2A1L | STON1-GTF2A1L readthrough | Regulation: Transcription | Ubiquitous |
| TAX1BP1 | Tax1 binding protein 1 | Immunity: Autophagy/NF-kB | Ubiquitous |
| TBX4 | T-box transcription factor 4 | Dev: Limb/Lung formation | Lung, Placenta, Prostate |
| TERF2IP | TERF2 interacting protein | Immunity/Telomere | Ubiquitous |
| TMC1 | Transmembrane channel like 1 | Auditory: Mechanotransduction | Inner Ear (Hair cells) |
| TMEM63C | Transmembrane protein 63C | Auditory: Osmosensitive channel | Brain, Pituitary gland |
| USP44 | Ubiquitin specific peptidase 44 | Ubiquitin: Deubiquitinase | Testis |
| USP8 | Ubiquitin specific peptidase 8 | Ubiquitin: Endosomal sorting | Ubiquitous |
| ZFHX4 | Zinc finger homeobox 4 | Neural: Differentiation | Ubiquitous |
| ZFP30 | ZFP30 zinc finger protein | Regulation: Transcription factor | Ubiquitous |
| ZNF778 | Zinc finger protein 778 | Regulation: Transcription factor | Ubiquitous |
| ZNF783 | Zinc finger protein 783 | Regulation: Transcription factor | Ubiquitous |

***Table S2:*** *Summary of 72 candidate genes identified by BUSTED-PH as exhibiting episodic diversifying selection (EDS) significantly associated with the evolution of echolocation in mammals (FDR $< 0.05$). For each gene, columns provide the official symbol, common gene name, primary biological function, and tissue expression profile derived from the Human Protein Atlas (HPA) RNA consensus dataset. Inclusion criteria required simultaneous evidence of positive selection on foreground branches and a significant selective regime difference between foreground and background lineages.*
