## Supplemental Table 3 for "BUSTED-PH: Isolating the genomic signatures of convergent phenotypes"

| Symbol | Gene | Function | Expression |
|---|---|---|---|
| ADAM21 | ADAM metallopeptidase domain 21 | Proteolysis: Metallopeptidase | Testis |
| ADRM1 | ADRM1 26S proteasome ubiquitin receptor | Ubiquitin: Proteasome receptor | Skeletal muscle |
| ARMCX6 | Armadillo repeat containing X-linked 6 | Unknown: Armadillo repeat | Ubiquitous |
| ASAP3 | ArfGAP with SH3 domain, ankyrin repeat and PH domain 3 | Signaling: ArfGAP | Ubiquitous |
| C6ORF132 | Chromosome 6 open reading frame 132 | Cilia: Unknown function | Esophagus, Skin 1 |
| CAPN12 | Calpain 12 | Proteolysis: Calpain | Liver, Gallbladder |
| CAPN15 | Calpain 15 | Proteolysis: Calpain | Ubiquitous |
| CCM2 | CCM2 scaffold protein | Dev: Vascular scaffold | Brain |
| CENPV | Centromere protein V | Cell cycle: Centromere | Ubiquitous |
| CUL7 | Cullin 7 | Growth: Ubiquitin ligase (3M) | Ubiquitous |
| DENND4B | DENN domain containing 4B | Transport: Rab GEF | Ubiquitous |
| DHX40 | DEAH-box helicase 40 | RNA: Helicase | Ubiquitous |
| DNAH11 | Dynein axonemal heavy chain 11 | Cilia: Dynein motor | Parathyroid gland |
| EMILIN2 | Elastin microfibril interfacer 2 | ECM: Elastin microfibril | Parathyroid gland |
| ENTPD7 | Ectonucleoside triphosphate diphosphohydrolase 7 | Metabolism: Nucleotide | Intestine |
| EPC2 | Enhancer of polycomb homolog 2 | Regulation: Chromatin remodeling | Ubiquitous |
| EXD1 | EXD1 3'-5' exoribonuclease | RNA: Exonuclease | Testis |
| FADS3 | Fatty acid desaturase 3 | Metabolism: Fatty acid | Ubiquitous |
| FGD5 | FYVE, RhoGEF and PH domain containing 5 | Dev: Angiogenesis RhoGEF | Ubiquitous |
| GCLC | Glutamate-cysteine ligase catalytic subunit | Metabolism: Glutathione synthesis | Liver |
| GLB1L2 | Galactosidase beta 1 like 2 | Metabolism: Galactosidase | Retina |
| GP5 | Glycoprotein V platelet | Blood: Platelet glycoprotein | Lymphoid tissue |
| GRIA1 | Glutamate ionotropic receptor AMPA type subunit 1 | Neural: Glutamate receptor | Brain, Retina |
| GRIN3A | Glutamate ionotropic receptor NMDA type subunit 3A | Neural: NMDA receptor | Brain |
| GSTCD | Glutathione S-transferase C-terminal domain containing | Other | Ubiquitous |
| H4C7 | H4 clustered histone 7 | Other | Not detected (HPA) |
| HMCN2 | Hemicentin 2 | ECM: Hemicentin | Intestine, Skeletal muscle, Endometrium 1 |
| HRH3 | Histamine receptor H3 | Neural: Histamine receptor | Brain |
| IQSEC2 | IQ motif and Sec7 domain ArfGEF 2 | Neural: ArfGEF | Skeletal muscle |
| ITPRID1 | ITPR interacting domain containing 1 | Regulation: ITPR interaction | Salivary gland, Brain, Skin 1 |
| KARS1 | Lysyl-tRNA synthetase 1 | Translation: tRNA synthetase | Ubiquitous |
| KIF17 | Kinesin family member 17 | Transport: Kinesin | Testis |
| LAMA2 | Laminin subunit alpha 2 | ECM: Laminin alpha 2 | Placenta |
| LAMA5 | Laminin subunit alpha 5 | ECM: Laminin alpha 5 | Ubiquitous |
| MACO1 | Macoilin 1 | Unknown: Macoilin | Ubiquitous |
| MAN2B2 | Mannosidase alpha class 2B member 2 | Metabolism: Mannosidase | Ubiquitous |
| MLLT3 | MLLT3 super elongation complex subunit | Regulation: Transcription | Ubiquitous |
| MMP10 | Matrix metallopeptidase 10 | ECM: Matrix metallopeptidase | Endometrium 1 |
| MN1 | MN1 proto-oncogene, transcriptional regulator | Dev: Transcriptional regulator | Skeletal muscle, Blood vessel |
| MTSS2 | MTSS I-BAR domain containing 2 | Cytoskeleton: I-BAR domain | Brain |
| MYH13 | Myosin heavy chain 13 | Muscle: Myosin heavy chain | Stomach 1, Skeletal muscle |
| MYH4 | Myosin heavy chain 4 | Muscle: Myosin heavy chain | Skeletal muscle |
| NOVA2 | NOVA alternative splicing regulator 2 | Neural: Splicing regulator | Brain |
| NPHS2 | NPHS2 stomatin family member, podocin | Kidney: Podocyte slit diaphragm | Kidney |
| NPIPB13 | Nuclear pore complex interacting protein family, member B13 | Transport: Nuclear pore | Ubiquitous |
| NPIPB5 | Nuclear pore complex interacting protein family member B5 | Transport: Nuclear pore | Ubiquitous |
| NYNRIN | NYN domain and retroviral integrase containing | Unknown: NYN domain | Ubiquitous |
| OR13C3 | Olfactory receptor family 13 subfamily C member 3 | Smell: Olfactory receptor | Not detected (HPA) |
| OR13C8 | Olfactory receptor family 13 subfamily C member 8 | Smell: Olfactory receptor | Not detected (HPA) |

| Symbol | Gene | Function | Expression |
|---|---|---|---|
| OR2A12 | Olfactory receptor family 2 subfamily A member 12 | Smell: Olfactory receptor | Not detected (HPA) |
| OR2G3 | Olfactory receptor family 2 subfamily G member 3 | Smell: Olfactory receptor | Not detected (HPA) |
| OR4K5 | Olfactory receptor family 4 subfamily K member 5 | Smell: Olfactory receptor | Not detected (HPA) |
| OR5I1 | Olfactory receptor family 5 subfamily I member 1 | Smell: Olfactory receptor | Not detected (HPA) |
| OR5L1 | Olfactory receptor family 5 subfamily L member 1 | Smell: Olfactory receptor | Not detected (HPA) |
| OR8B2 | Olfactory receptor family 8 subfamily B member 2 | Smell: Olfactory receptor | Testis |
| PCDHB5 | Protocadherin beta 5 | Neural: Protocadherin | Ubiquitous |
| PCDHGA9 | Protocadherin gamma subfamily A, 9 | Neural: Protocadherin | Ubiquitous |
| PLAA | Phospholipase A2 activating protein | Ubiquitin: Degradation | Ubiquitous |
| PLK3 | Polo like kinase 3 | Cell cycle: Kinase | Ubiquitous |
| PRG4 | Proteoglycan 4 | ECM: Lubricin | Liver, Adipose tissue |
| RAP1GAP2 | RAP1 GTPase activating protein 2 | Signaling: GTPase activating | Pancreas, Bone marrow |
| RASIP1 | Ras interacting protein 1 | Dev: Vascular stability | Ubiquitous |
| RNH1 | Ribonuclease/angiogenin inhibitor 1 | RNA: Ribonuclease inhibitor | Ubiquitous |
| RSAD2 | Radical S-adenosyl methionine domain containing 2 | Immunity: Antiviral | Salivary gland |
| RUVBL1 | RuvB like AAA ATPase 1 | Regulation: Chromatin remodeling | Ubiquitous |
| SCARF2 | Scavenger receptor class F member 2 | Metabolism: Scavenger receptor | Blood vessel |
| SCN4B | Sodium voltage-gated channel beta subunit 4 | Ion channel: Sodium beta | Tongue, Skeletal muscle, Brain |
| SLC22A13 | Solute carrier family 22 member 13 | Transport: Solute carrier | Kidney |
| SMC1A | Structural maintenance of chromosomes 1A | Cell cycle: Cohesin | Ubiquitous |
| SOAT1 | Sterol O-acyltransferase 1 | Metabolism: Cholesterol | Adrenal gland |
| SRRM1 | Serine and arginine repetitive matrix 1 | Splicing: RNA processing | Ubiquitous |
| STK25 | Serine/threonine kinase 25 | Signaling: Kinase | Skeletal muscle |
| SYNE2 | Spectrin repeat containing nuclear envelope protein 2 | Other | Skeletal muscle |
| TAF4 | TATA-box binding protein associated factor 4 | Regulation: Transcription | Ubiquitous |
| TAX1BP1 | Tax1 binding protein 1 | Immunity: Autophagy/NF-kB | Ubiquitous |
| TBC1D16 | TBC1D16, TBC1 domain family member 16 | Transport: Rab GAP | Ubiquitous |
| TDRD12 | Tudor domain containing 12 | Sperm: Piwi pathway | Testis |
| TERF2IP | TERF2 interacting protein | Other | Ubiquitous |
| TMEM63C | Transmembrane protein 63C | Other | Brain, Pituitary gland |
| TMEM94 | Transmembrane protein 94 | Other | Ubiquitous |
| TNC | Tenascin C | ECM: Extracellular matrix | Smooth muscle, Blood vessel |
| TOR1AIP2 | Torsin 1A interacting protein 2 | Other | Ubiquitous |
| TTLL5 | Tubulin tyrosine ligase like 5 | Other | Testis |
| TUBGCP6 | Tubulin gamma complex associated protein 6 | Microtubule: Nucleation | Ubiquitous |
| WWC3 | WWC family member 3 | Other | Ubiquitous |
| ZMYM3 | Zinc finger MYM-type containing 3 | Other | Ubiquitous |
| ZNF282 | Zinc finger protein 282 | Other | Ubiquitous |
| ZNF598 | Zinc finger protein 598, E3 ubiquitin ligase | Other | Ubiquitous |
| ZNF777 | Zinc finger protein 777 | Other | Ubiquitous |
| ZNF853 | Zinc finger protein 853 | Other | Ubiquitous |
| ZSCAN10 | Zinc finger and SCAN domain containing 10 | Other | Not detected (HPA) |

***Table S3:*** *Summary of 91 candidate genes identified by BUSTED-PH as exhibiting episodic diversifying selection (EDS) significantly associated with the evolution of large body size in mammals (FDR < 0.05). For each gene, columns provide the official symbol, common gene name, primary biological function, and tissue expression profile derived from the Human Protein Atlas (HPA) RNA consensus dataset. Inclusion criteria required simultaneous evidence of positive selection on foreground branches and a significant selective regime difference between foreground and background lineages.*
