## Supplemental Table 4 for "BUSTED-PH: Isolating the genomic signatures of convergent phenotypes"

| Method | Approach | Pros | Cons | Assumptions | Compute Load |
|---|---|---|---|---|---|
| **BUSTED-PH** | Branch-site codon model contrasting foreground vs. background selection regimes. | Explicitly tests for trait-associated selection; controls for background selection; detects gene-wide episodic selection; robust to some alignment error. | Requires a tree and alignment; gene-level resolution (may miss single-site drivers). | Independent codon sites; random effects dN/dS distribution; shared gene tree. | Moderate (ML) |
| **Standard Branch-Site (e.g., PAML)** | Tests for positive selection on foreground branches against a neutral null. | Well-established; widely used; sensitive to episodic selection on specified lineages. | Cannot distinguish trait-associated selection from pervasive background selection; high FP rate if background is adaptive. | Foreground specified a priori; background evolves neutrally/conservatively. | Moderate-High |
| **RERConverge [Kowalczyk et al., 2019]** | Correlates relative evolutionary rates (RER) of genes with phenotype across a phylogeny. | Fast genome-wide scans; detects shifts in constraint (e.g., relaxed selection); accounts for tree structure. | Low resolution (gene-wide rate only); cannot detect site-specific positive selection (dN/dS). | Rate variation reflects functional constraint; reliable branch length estimates. | Low |
| **PhyloAcc [Hu et al., 2019]** | Bayesian model detecting acceleration of substitution rates in conserved elements. | Designed for non-coding elements; handles missing data well; detects lineage-specific acceleration. | Focuses on overall rate acceleration (often loss/relaxation) rather than adaptive protein evolution. | Acceleration implies function loss or gain; conserved elements are comparable. | High (MCMC) |
| **Forward Genomics [Prudent et al., 2016]** | Assesses correlation between phenotype presence/loss and sequence divergence/loss. | Good for detecting "use it or lose it" gene loss associated with trait loss. | Low power for detecting adaptive gain of function; sensitive to phylogenetic artifacts. | Trait loss leads to relaxation/loss of functional genes. | Low-Moderate |
| **Topology Tests / SSLS [Parker et al., 2013]** | Compares likelihood support for gene alignments under species tree vs. convergent topologies. | Detects cumulative convergent signal sufficient to override species history. | Sensitive to ILS/hemiplasy; requires defining specific alternative topologies; computationally expensive for many trees. | Convergent substitutions drive gene tree toward phenotypic grouping. | High |
| **CSUBST [Fukushima and Pollock, 2023]** | Metric ($\omega_C$) quantifying correlation of site-specific amino acid preferences with phenotype. | Detects specific convergent amino acid substitutions; explicitly corrects for error. | Requires identical/similar substitutions; limited by "Stokes Shift" (entrenchment). | Convergence manifests as identical/similar amino acid shifts. | High |
| **TraitRELAX [Halabi et al., 2021]** | Codon model testing for relaxation of selection (k parameter) associated with a trait. | Specifically models relaxed selection (trait loss); distinguishes relaxation from positive selection. | Primarily tests for relaxation, not positive selection. | Selection relaxation ($k < 1$) correlates with trait. | Moderate (ML) |
| **ACEP [Cao et al., 2025]** | Protein language model embeddings to detect convergent evolutionary patterns. | Captures high-order dependencies and physicochemical properties beyond simple substitution. | Interpretation can be difficult (black box); requires significance testing framework. | PLM embeddings capture functional equivalence; convergence is detectable in latent space. | High (Inference) |
| **Sparse Learning [Allard et al., 2025]** | Lasso-based sparse learning to identify specific sites driving phenotype-genotype association. | Feature selection identifies specific codon drivers; handles high-dimensional data. | Requires large sample sizes for stability; sparse assumption may not hold for all traits. | Adaptive substitutions are sparse; linear relationship between sites and phenotype. | Moderate-High |

***Table S4:*** *Head-to-head comparison of* `BUSTED-PH` *with other comparative genomic methods for detecting trait-associated evolution.* `BUSTED-PH` *offers a distinct advantage by explicitly modeling and contrasting selective regimes (dN/dS) between foreground and background, filtering out pervasive selection that confounds standard branch-site models and offering higher resolution than rate-based approaches.*
